# Optical determination of absolute membrane potential

**DOI:** 10.1101/519736

**Authors:** Julia R. Lazzari-Dean, Anneliese M.M. Gest, Evan W. Miller

## Abstract

All cells maintain ionic gradients across their plasma membranes, producing transmembrane potentials (Vmem). Mounting evidence suggests a relationship between resting Vmem and the physiology of non-excitable cells with implications in diverse areas, including cancer, cellular differentiation, and body patterning. A lack of non-invasive methods to record absolute Vmem limits our understanding of this fundamental signal. To address this need, we developed a fluorescence lifetime-based approach (VF-FLIM) to visualize and optically quantify Vmem with single-cell resolution. Using VF-FLIM, we report Vmem distributions over thousands of cells, a 100-fold improvement relative to electrophysiological approaches. In human carcinoma cells, we visualize the voltage response to epidermal growth factor stimulation, stably recording a 10-15 mV hyperpolarization over minutes. Using pharmacological inhibitors, we identify the source of the hyperpolarization as the Ca^2+^-activated K^+^ channel K_ca_3.1. The ability to optically quantify absolute Vmem with cellular resolution will allow a re-examination of its roles as a cellular signal.

## Introduction

Membrane potential (V_mem_) is an essential facet of cellular physiology. In electrically excitable cells, such as neurons and cardiomyocytes, voltage-gated ion channels enable rapid changes in membrane potential. These fast membrane potential changes, on the order of milliseconds to seconds, trigger release of neurotransmitters in neurons or contraction in myocytes. The resting membrane potential of these cells, which changes over longer timescales, affects their excitability. In non-electrically excitable cells, slower changes in V_mem_—on the order of seconds to hours—are linked to a variety of fundamental cellular processes^1^, including mitosis ^2^, cell cycle progression ^3^, and differentiation ^4^ At the tissue and organismal level, mounting lines of evidence point to the importance of electrochemical gradients in development, body patterning, and regeneration ^5^.

Despite the importance of membrane potential to diverse processes over a range of time scales, the existing methods for recording V_mem_ are inadequate for characterizing distributions of V_mem_ states in a sample or studying gradual shifts in resting membrane potential (**Figure 1- supplement 1**). Patch clamp electrophysiology remains the gold standard for recording cellular electrical parameters, but it is low throughput, highly invasive, and difficult to implement over extended time periods. Where reduced invasiveness or higher throughput analyses of V_mem_ are required, optical methods for detecting events involving V_mem_ changes (e.g. whether an action potential occurred) are often employed ^6–8^. However, optical approaches generally use fluorescence intensity values as a readout, which cannot report either the absolute values of V_mem_ or the absolute amount by which V_mem_ changed ^9^. Variations in dye loading, illumination intensity, fluorophore bleaching, and/or cellular morphology dramatically complicate fluorescence intensity measurements, making calibration and determination of actual membrane potential difficult or impossible. This limitation restricts optical analysis to detection of acute V_mem_ changes, which can be analyzed without comparisons of V_mem_ between cells or over long timescales. Two-component systems, with independent wavelengths for ratio-based calibration, have seen limited success ^10^, and they confer significant capacitive load on the cell ^11^. Further, their performance hinges on carefully tuned loading procedures of multiple lipophilic indicators ^12^, which can be challenging to reproduce across different samples and days.

To quantify a parameter such as voltage or concentration from a single-color fluorescence signal, fluorescence lifetime (τ_fl_) imaging (FLIM) can be employed instead of conventional fluorescence microscopy. By measuring the fluorescence lifetime, an intrinsic property of the sensor, FLIM avoids many of the artifacts that confound extrinsic fluorescence intensity measurements. As a result, FLIM can be calibrated to reproducibly and quantitatively report biological properties if the analyte or property in question affects the lifetime of the probe’s fluorescent excited state. FLIM has been successfully employed to record a number of biochemical and biophysical parameters, including intracellular Ca^2+^ concentration ^13^, viscosity ^14^, GTPase activity ^15^, kinase activity ^16^, and redox state (NADH/NAD+ ratio)^17^, among others ^18^. Attempts to record absolute voltage with FLIM, however, have been limited in success ^19–21^. Previous work focused on genetically-encoded voltage indicators (GEVIs), which have complex relationships between τfl and voltage ^20^ and low sensitivity to voltage in lifetime ^21^. Because of their poor voltage resolution, the fluorescence lifetimes of these GEVIs cannot be used to detect most biologically relevant voltage changes, which are on the order of tens of millivolts.

Fluorescent voltage indicators that use photoinduced electron transfer (PeT) as a voltage-sensing mechanism are promising candidates for a FLIM-based approach to optical V_mem_ quantification. Because PeT affects the nonradiative decay rate of the fluorophore excited state, it has been successfully translated from intensity to τfl imaging with a number of small molecule probes for Ca^2+^ ^22^. We previously established that VoltageFluor (VF)-type dyes transduce changes in cellular membrane potential to changes in fluorescence intensity and that the voltage response of VF dyes is consistent with a photoinduced electron transfer (PeT)-based response mechanism ^23,24^ Changes in the transmembrane potential alter the rate of PeT ^25,26^ from an electron-rich aniline donor to a fluorescent reporter, thereby modulating the fluorescence intensity of VF dyes ^23^ (**Fig. 1A,B**). VoltageFluors also display low toxicity and rapid, linear responses to voltage.

**Fig. 1.**
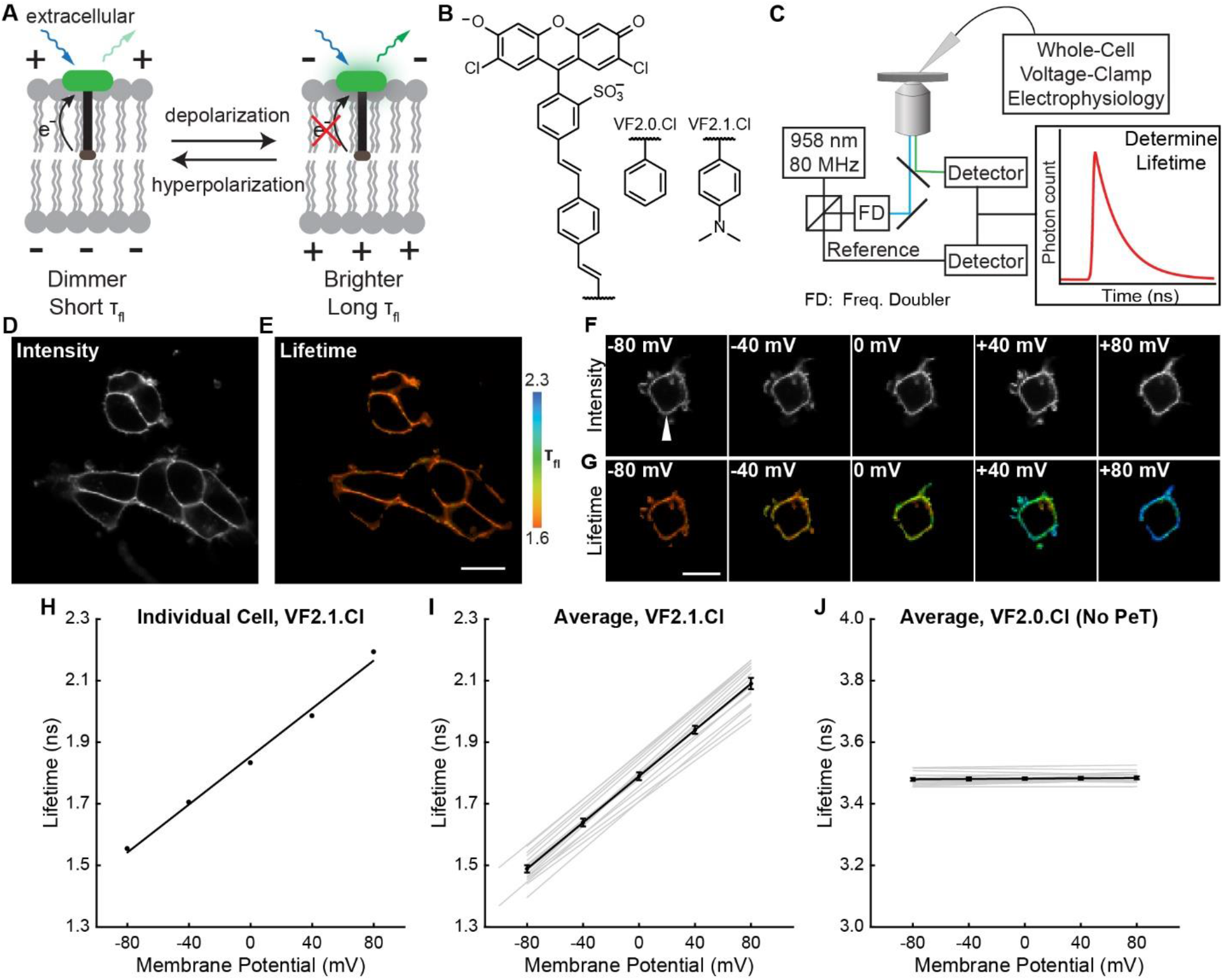
VoltageFluor FLIM linearly reports absolute membrane potential. (**A**) Mechanism of VoltageFluor dyes, in which depolarization of the membrane potential attenuates the rate of photoinduced electron transfer. (**B**) Structures of the VF molecules used in this study. (**C**) Schematic of the TCSPC system used to measure fluorescence lifetime. Simultaneous electrophysiology was used to establish lifetime-voltage relationships. (**D**) Fluorescence intensity and (**E**) lifetime of HEK293T cells loaded with 100 nM VF2.1.Cl. (**F**) Intensity and (**G**) lifetime images of HEK293T cells voltage clamped at the indicated membrane potential. (**H**) Quantification of the single trial shown in (**G**), with a linear fit to the data. (**I**) Evaluation of VF2.1.Cl lifetime-voltage relationships in many individual HEK293T cells. Gray lines represent linear fits on individual cells. Black line is the average lifetime-voltage relationship across all cells (n=17). (**J**) VF2.0.Cl lifetime does not exhibit voltage-dependent changes. Gray lines represent linear fits on individual cells, and the black line is the average lifetime-voltage relationship across all cells (n=17). Scale bars represent 20 μm. Error bars represent mean ± SEM.

Here, we develop fluorescence lifetime imaging of VoltageFluor dyes (VF-FLIM) as a quantitative, all-optical approach for recording absolute membrane potential with single cell resolution. Using patch-clamp electrophysiology as a standard, we demonstrate that the fluorescence lifetime of the VoltageFluor dye VF2.1.Cl reports absolute membrane potential with >20-fold improved accuracy over previous optical approaches. To highlight the 100-fold increase in throughput over manual patch-clamp electrophysiology, we record resting membrane potentials of thousands of cells. To our knowledge, this work represents the first broad view of the distribution of resting membrane potentials present *in situ*. To showcase the spatiotemporal and voltage resolution of VF-FLIM, we quantify the gradual, small voltage changes that arise from growth factor stimulation of human carcinoma cells. Through pharmacological perturbations, we conclude that the voltage changes following epidermal growth factor (EGF) stimulation arise from activation of the calcium-activated potassium channel Koa3.1. Our results show that fluorescence lifetime of VF dyes is a generalizable and effective approach for studying resting membrane potential in a range of biological contexts.

## Results

### VoltageFluor Fluorescence Lifetime Varies Linearly with Membrane Potential

To characterize how the photoinduced electron transfer process affects fluorescence lifetime, we compared the τfl of the voltage-sensitive dye VF2.1.Cl with its voltage-insensitive counterpart VF2.0.Cl (**Fig. 1B**). We recorded the τfl of bath-applied VF dyes in HEK293T cells using time-correlated single-photon counting (TCSPC) FLIM (**Fig. 1C-E**). VF2.1.Cl is localized to the plasma membrane and exhibits a biexponential τfl decay with decay constants of approximately 0.9 and 2.6 ns (**Scheme S2**). For all subsequent analysis of VF2.1.Cl lifetime, we refer to the weighted average τfl, which is approximately 1.6 ns in HEK293T cell membranes at rest. VF2.0.Cl (**Fig. 1B**), which lacks the aniline substitution and is therefore voltage-insensitive ^24^, shows a τfl of 3.5 ns in cell membranes, which is similar to the lifetime of an unsubstituted fluorescein ^27^ (**Fig. 1-supplement 2**). We also examined VoltageFluor lifetimes at a variety of dye loading concentrations to test for concentration-dependent changes in dye lifetime, which have been reported for fluorescein derivatives ^28^. Shortened VF lifetimes were observed at high dye concentrations (**Fig. 1-supplement 3**); all subsequent VF-FLIM studies were conducted at dye concentrations low enough to avoid this concentration-dependent change in lifetime.

To assess the voltage dependence of VoltageFluor τfl, we controlled the plasma membrane potential of HEK293T cells with whole-cell voltage-clamp electrophysiology while simultaneously measuring the τfl of VF2.1.Cl (**Fig. 1C**). Single-cell recordings show a linear τfl response to applied voltage steps, and individual measurements deviate minimally from the linear fit (**Fig. 1F-H**). VF2.1.Cl τfl is reproducible across different cells at the same resting membrane potential, allowing determination of V_mem_ from τfl images taken without concurrent electrophysiology (**Fig. 1I**). Voltage-insensitive VF2.0.Cl shows no τfl change in response to voltage (**Fig. 1J, Fig. 1-supplement 4**), consistent with a τfl change in VF2.1.Cl arising from a voltage-dependent PeT process. In HEK293T cells, VF2.1.Cl exhibits a sensitivity of 3.50 ± 0.08 ps/mV and a 0 mV lifetime of 1.77 ± 0.02 ns, corresponding to a fractional change in τfl (Δτ/τ) of 22.4 ± 0.4% per 100 mV. These values are in good agreement with the 27% ΔF/F intensity change per 100 mV originally observed for VF2.1.Cl ^23,24^ To estimate the voltage resolution of VF-FLIM, we analyzed the variability in successive measurements on the same cell (intra-cell resolution) and on different cells (inter-cell resolution, see **Methods**). We estimate that the resolution for tracking and quantifying voltage changes in a single HEK293T cell is 4 mV (intra-cell resolution), whereas the resolution for single-trial determination of a particular HEK293T cell’s absolute V_mem_ is 20 mV (inter-cell resolution).

We compared the performance of VF-FLIM to that of CAESR, the best previously reported GEVI for optically recording absolute membrane potential using FLIM ^21^. Using simultaneous FLIM and voltage-clamp electrophysiology, we determined the relationship between τfl and V_mem_ for the genetically encoded voltage indicator CAESR under 1 photon excitation (**Fig. 1- supplement 5**). We recorded a sensitivity of −1.2 ± 0.1 ps/mV and a 0 mV lifetime of 2.0 ± 0.2 ns, which corresponds to a −6.1 ± 0.8% Δτ/τ per 100 mV (mean ± SEM of 9 measurements), in agreement with the reported sensitivity of −0.9 ps/mV and 0 mV lifetime of 2.7 ns with 2 photon excitation ^21^. Relative to VF2.1.Cl, CAESR displays 3-fold lower sensitivity (−1.2 ps/mV vs 3.5 ps/mV in HEK293T cells) and 7-fold higher voltage-independent variability in lifetime (0.46 ns vs 0.07 ns, standard deviation of the 0 mV lifetime measurement). For CAESR in HEK293T cells, we calculate a voltage resolution of 37 ± 7 mV for quantifying voltage changes on an individual cell (intra-cell, compared to 4 mV for VF2.1.Cl, see **Methods**) and resolution of 390 mV for determination of a particular cell’s absolute V_mem_ (inter-cell, compared to 20 mV for VF2.1.Cl). Because cellular resting membrane potentials and voltage changes (e.g. action potentials) are on the order of tens of millivolts, VF-FLIM has sufficient resolution for biologically relevant V_mem_ recordings, whereas CAESR does not.

### Evaluation of VF-FLIM across Cell Lines and Culture Conditions

The voltage-dependent τfl response of VF2.1.Cl is generalizable across different cell types. We calibrated VF-FLIM in four additional commonly used cell lines: A431, CHO, MDA-MB-231, and MCF-7 (**Fig. 2, Fig. 2-supplement 1, Fig. 2- supplement 2**). All cells displayed a linear relationship between VF τfl and V_mem_, with average sensitivities of 3.1 to 3.7 ps/mV and average 0 mV lifetimes ranging from 1.74 to 1.87 ns. In all cases, we observed better voltage resolution for quantification of V_mem_ changes on a given cell versus comparisons of absolute V_mem_ between cells. For all cell lines tested, the changes in voltage for a given cell could be quantified with resolutions at or better than 5 mV (intra-cell resolution, Methods). For absolute V_mem_ determination of a single cell, we observed voltage resolutions ranging from 11 to 24 mV (intercell resolution, **Fig. 2-supplement 3**). The inter-cell resolution of VF-FLIM appears to be cell-type dependent; MCF-7 cells displayed greater variability than other cell lines tested (**Fig. 2B, Fig. 2-supplement 3**).

**Fig. 2.**
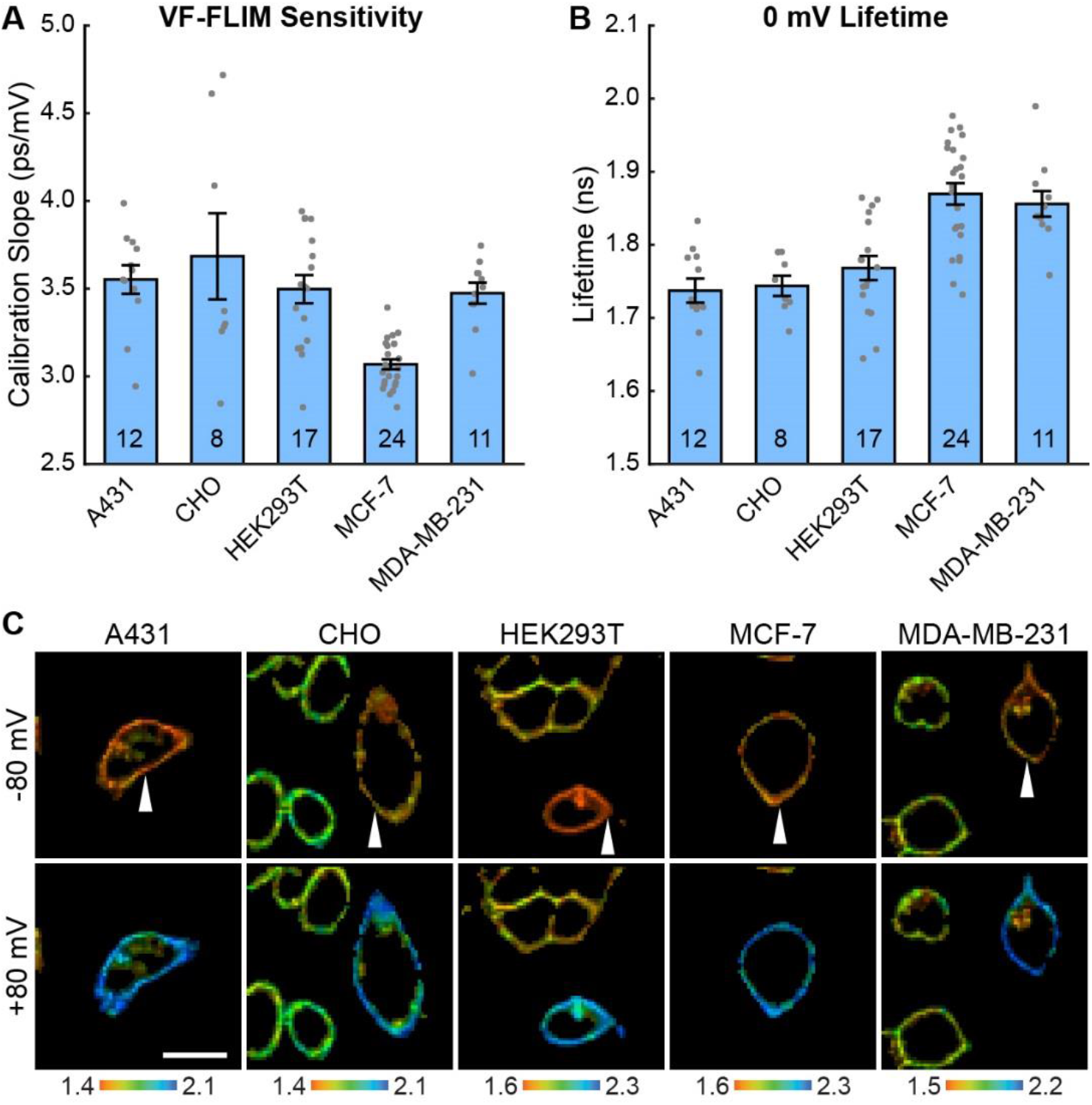
VF-FLIM is a general and portable method for optically determining membrane potential. VF2.1.Cl lifetime-voltage relationships were determined with whole cell voltage clamp electrophysiology in five cell lines. (**A**) Slope and (**B**) 0 mV reference point of linear fits for the lifetime-voltage relationship, shown as mean ± S.E.M. Gray dots are single cells. (**C**) Representative lifetime-intensity overlay images for each cell line with the indicated cells (white arrow) held at −80 mV (top) or +80 mV (bottom). Lifetime scales are in ns. Scale bar is 20 μm.

To verify that VF-FLIM was robust in groups of cells in addition to the isolated, single cells generally used for patch clamp electrophysiology, we determined lifetime-voltage relationships for small groups of A431 cells (**Fig. 2-supplement 4A-E**). We found that calibrations made in small groups of cells are nearly identical to those obtained on individual cells, indicating that VF-FLIM only needs to be calibrated once for a given type of cell. For pairs or groups of three cells we recorded a sensitivity of 3.3 ± 0.2 ps/mV and a 0 mV lifetime of 1.78 ± 0.02 ns (mean ± SEM of 5 pairs and 2 groups of 3; values are for the entire group, not just the cell in contact with the electrode), which is similar to the sensitivity of 3.55 ± 0.08 ps/mV and 0 mV lifetime of 1.74 ± 0.02 ns we observe in single A431 cells. The slight reduction in sensitivity seen in cell groups is likely attributable to space clamp error, which prevents complete voltage clamp of the cell group ^29,30^. Indeed, when we analyzed only the most responsive cell in the group (in contact with the electrode), we obtained a slope of 3.7 ± 0.1 ps/mV and 0 mV lifetime of 1.79 ± 0.02 ns, in good agreement with the single cell data. The space clamp error can be clearly visualized (**Figure 2 – supplement 4E**), where one cell in the group of 3 responded much less to the voltage command.

To test whether VF-FLIM is also extensible to cells maintained with different culture conditions, we recorded lifetime-V_mem_ relationship in serum-starved A431 cells (**Figure 2 – supplement 4F-K**), obtaining an average sensitivity of 3.6 ± 0.1 ps/mV and a 0 mV lifetime of 1.76 ± 0.01 ns (n=2 single cells, 2 pairs, 3 groups of 3 cells), in excellent agreement with the values obtained for non-serum starved cells. We also tested for concentration-dependent changes in VF lifetime in all five cell lines and in serum starvation conditions. Similar to VF2.1.Cl lifetime in HEK293T cells (**Fig. 1-supplement 3**), we observed shortening of VF2.1.Cl lifetimes between 200 and 500 nM dye in all cases (**Figure 2-supplement 5**). All subsequent experiments were carried out at VF2.1.Cl concentrations well below the regime where VF concentration-dependent lifetime changes were observed.

### Optical Determination of Resting Membrane Potential Distributions

The throughput of VF-FLIM enables cataloging of resting membrane potentials of thousands of cells in only a few hours of the experimenter’s time. We optically recorded resting membrane potential distributions for A431, CHO, HEK293T, MCF-7, and MDA-MB-231 cells using VF-FLIM (**Fig. 3, Fig. 3 – supplement 1, Fig. 3 – supplement 2**). We report resting membrane potentials by cell group (**Methods, Scheme S2**) because adjacent cells in these cultures are electrically coupled to some degree via gap junctions ^31^. Each group of cells represents an independent sample for V_mem_. In addition, the fluorescent signal originating from membranes of adjacent cells cannot be separated with a conventional optical microscope, so assignment of a region of membrane connecting multiple cells would be arbitrary. VF-FLIM images (**Fig. 3, Fig. 3 – supplement 1, Fig. 3 - supplement 2**) contain spatially resolved voltage information, but caution should be employed in interpreting pixel to pixel differences in lifetime. Because VF-FLIM was calibrated here using the average plasma membrane τfl for each cell, optical V_mem_ should be interpreted per cell or cell group.

**Fig. 3.**
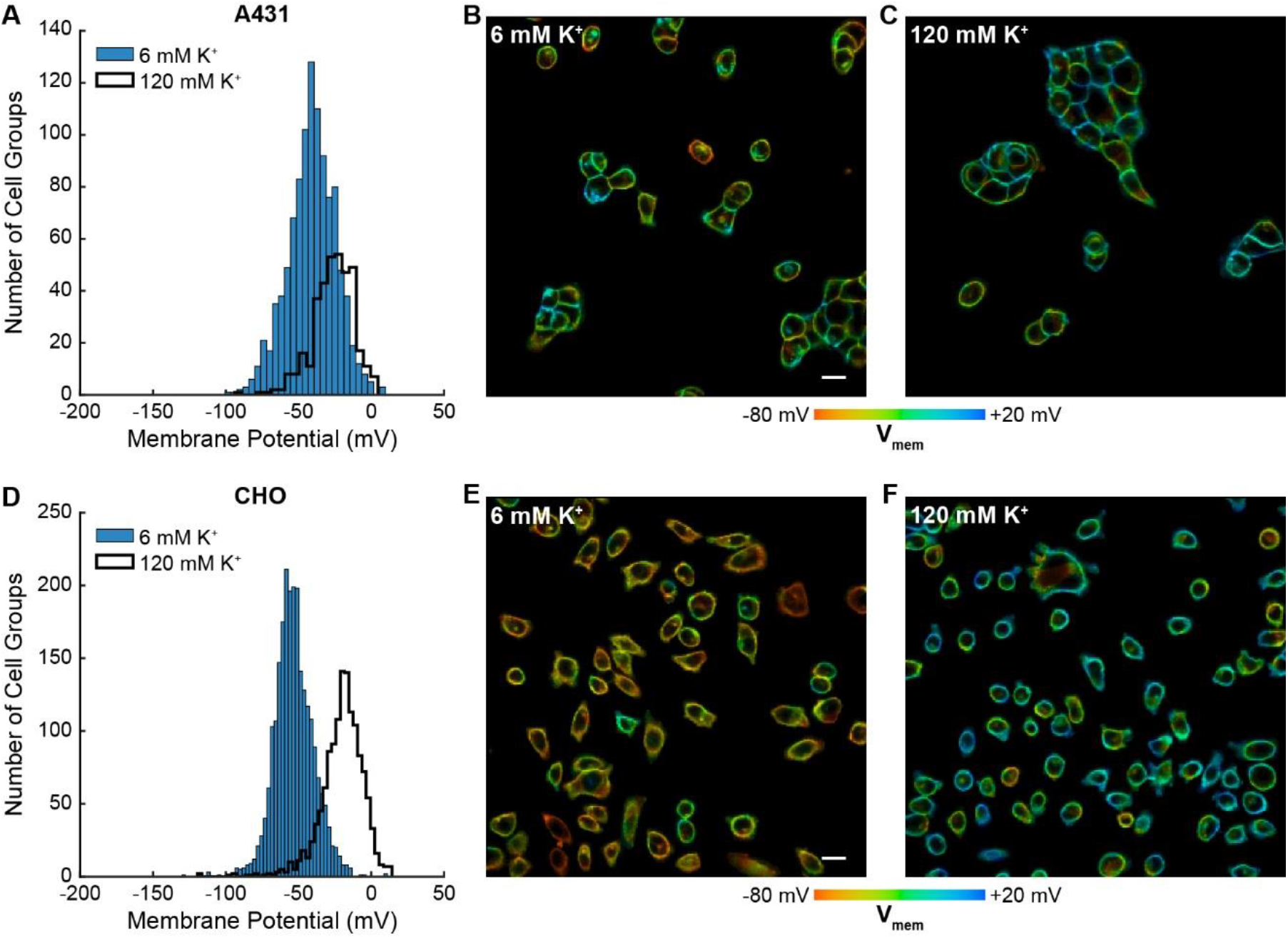
Rapid optical profiling of V_mem_ at rest and in high extracellular K^+^. Fluorescence lifetime images of cells incubated with 100 nM VF2.1.Cl were used to determine V_mem_ from previously performed electrophysiological calibration (**Fig. 2**). (**A**) Histograms of V_mem_ values recorded in A431 cells incubated with 6 mM extracellular K^+^ (commercial HBSS, n=1056) or 120 mM K^+^ (high K^+^ HBSS, n=368). (**B**) Representative lifetime image of A431 cells in 6 mM extracellular K^+^. (**C**) Representative lifetime image of A431 cells in 120 mM extracellular K^+^. (**D**) Histograms of V_mem_ values observed in CHO cells under normal (n=2410) and high K^+^ (n=1310) conditions. Representative lifetime image of CHO cells in (**E**) 6 mM and (**F**) 120 mM extracellular K^+^. Bin sizes were determined by the Freedman-Diaconis rule. Intensities in the lifetime-intensity overlay images are not scaled to each other. Scale bars, 20 μm.

Mean resting membrane potentials recorded by VF-FLIM range from −53 to −29 mV, depending on the cell line. These average V_mem_ values fall within the range reported in the literature for all of the cell lines we measured (**Fig. 3 - supplement 3**). We also recorded resting membrane potentials in a high K^+^ buffer (120 mM K^+^, “high K^+^ HBSS”), where we observed a depolarization of 15 to 41 mV, bringing the mean V_mem_ up to −26 mV to +4 mV, again depending on the cell line. Our optical determination of V_mem_ is in good agreement with theory: the Goldman-Hodgkin-Katz equation ^32^ predicts V_mem_ of −91 to −27 mV in 6 mM extracellular K^+^ and −25 to +2 mV in 120 mM extracellular K^+^, depending on ion permeability and intracellular ion concentration (see **Methods**).

### Membrane potential dynamics in epidermal growth factor signaling

We thought VF-FLIM was a promising method for elucidating the roles of membrane potential in non-excitable cell signaling. Specifically, we wondered whether VF-FLIM might be well-suited to dissect conflicting reports surrounding changes in membrane potential during EGF/EGF receptor (EGFR)-mediated signaling. Receptor tyrosine kinase (RTK)-mediated signaling is a canonical signaling paradigm for eukaryotic cells, transducing extracellular signals into changes in cellular state. Although the involvement of second messengers like Ca^2+^, cyclic nucleotides, and lipids are well characterized, membrane potential dynamics and their associated roles in non-excitable cell signaling remain less well-defined. In particular, the activation of EGFR via EGF has variously been reported to be depolarizing ^33^, hyperpolarizing ^34^, or electrically silent ^35,36^

We find that treatment of A431 cells with EGF results in a 15 mV hyperpolarization within 60-90 seconds in approximately 80% of cells (**Fig. 4A-C, Fig. 4-supplement 1, Fig. 4 – supplement 2**), followed by a slow return to baseline within 15 minutes (**Fig. 4D-F, Fig. 4-supplement 3**). The voltage response to EGF is dose-dependent, with an EC50 of 90 ng/mL (14 nM) (**Fig. 4-supplement 4**). Vehicle-treated cells show very little τfl change (**Fig. 4A-F**). Identical experiments with voltage-insensitive VF2.0.Cl (**Fig. 4G-H, Fig. 4 – supplement 1, Fig. 4 – supplement 3, Fig. 4 – supplement 5**) reveal little change in τfl upon EGF treatment, indicating the drop in τfl arises from membrane hyperpolarization. We observe the greatest hyperpolarization 1 to 3 minutes after treatment with EGF, which is abolished by inhibition of EGFR and ErbB2 tyrosine kinase activity with the covalent inhibitor canertinib (**Fig. 4I-J, Fig. 4-supplement 6**). Blockade of the EGFR kinase domain with gefitinib, a non-covalent inhibitor of EGFR, also results in a substantial decrease in the EGF-evoked hyperpolarization (**Fig. 4I-J, Fig. 4-supplement 6**). Together, these results indicate that A431 cells exhibit an EGF-induced hyperpolarization, which depends on the kinase activity of EGFR and persists on the timescale of minutes.

**Fig. 4.**
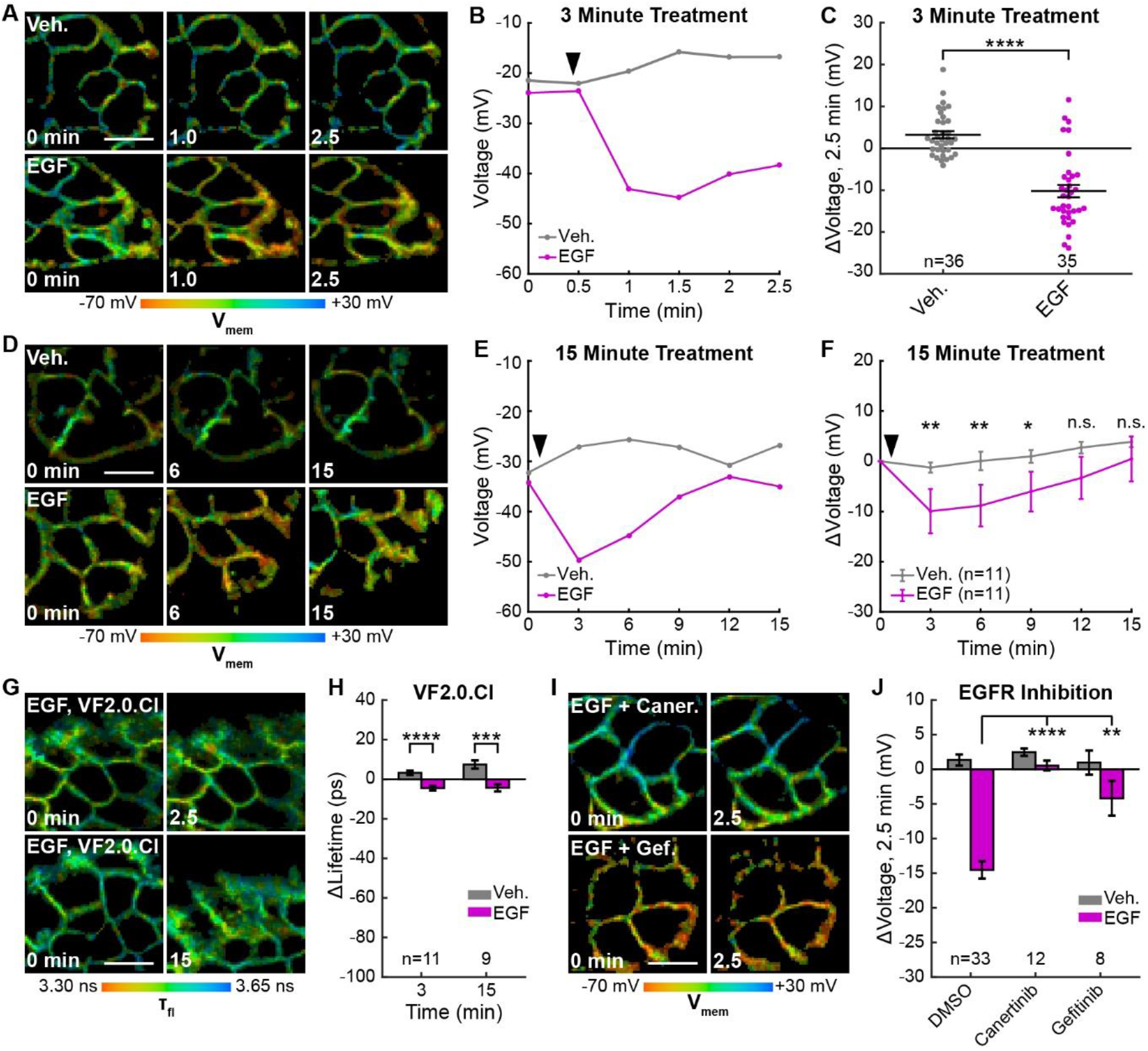
EGFR-mediated receptor tyrosine kinase activity produces a transient hyperpolarization in A431 cells. (**A**) Representative VF-FLIM time series of A431 cells treated with imaging buffer vehicle (top) or 500 ng/mL EGF (80 nM, bottom). (**B**) Quantification of images in (**A**), with Vehicle (Veh.)/EGF added at black arrow. (**C**) Aggregated responses for various trials of cells treated with vehicle or EGF. (**D**) Lifetime images of longer-term effects of vehicle (top) or EGF (bottom) treatment. (**E**) Quantification of images in (**D**). (**F**) Average response of cells over the longer time course. (**G**) Images of VF2.0.Cl (voltage insensitive) lifetime before and after EGF treatment. No τfl treatment. No τfl change is observed 2.5 (top) or 15 minutes (bottom) following EGF treatment. (**H**) Average VF2.0.Cl lifetime changes following EGF treatment. VF2.0.Cl graphs and images are scaled across the same lifetime range (350 ps) as VF2.1.Cl plots and images. The small drift observed would correspond to 2-4 mV of voltage change in VF2.1.Cl lifetime. (**I**) Lifetime images of A431 cells before and after EGF addition, with 500 nM canertinib (top) or 10 μM gefitinib (bottom). (**J**) Voltage changes 2.5 minutes after EGF addition in cells treated with DMSO (vehicle control) or an EGFR inhibitor. Scale bars are 20 μm. (C,F,H): Asterisks indicate significant differences between vehicle and EGF at that time point. (J): Asterisks reflect significant differences between EGF-induced voltage responses with DMSO vehicle or an EGFR inhibitor (n.s. p>0.05, * p<0.05, ** p<0.01, *** p<0.001, **** p<0.0001, two-tailed, unpaired, unequal variances t-test).

Outward K^+^ currents could mediate EGF-induced hyperpolarization. Consistent with this hypothesis, dissipation of the K^+^ driving force by raising extracellular [K^+^] completely abolishes the typical hyperpolarizing response to EGF and instead results in a small depolarizing potential of approximately 3 mV (**Fig. 5A, Fig. 5 – supplement 1B**). Blockade of voltage-gated K^+^ channels (Kv) with 4-aminopyridine (4-AP) prior to EGF treatment enhances the hyperpolarizing response to EGF (**Fig. 5A, 5B, Fig. 5-supplement 1C**). In contrast, blockade of Ca^2+^-activated K^+^ channels (Koa) with charybdotoxin (CTX) results in a depolarizing potential of approximately 4 mV after exposure to EGF, similar to that observed with high extracellular [K^+^] (**Fig. 5A, 5B, Fig. 5- supplement 1D**). TRAM-34, a specific inhibitor of the intermediate-conductance Ca^2+^ activated potassium channel Kea3.1 ^37^, also abolishes EGF-induced hyperpolarization (**Fig. 5A, Fig. 5- supplement 1E**). CTX treatment has little effect on the resting membrane potential, while TRAM-34 or 4-AP depolarizes cells by approximately 5-10 mV (**Fig. 5-supplement 2**).

**Fig. 5.**
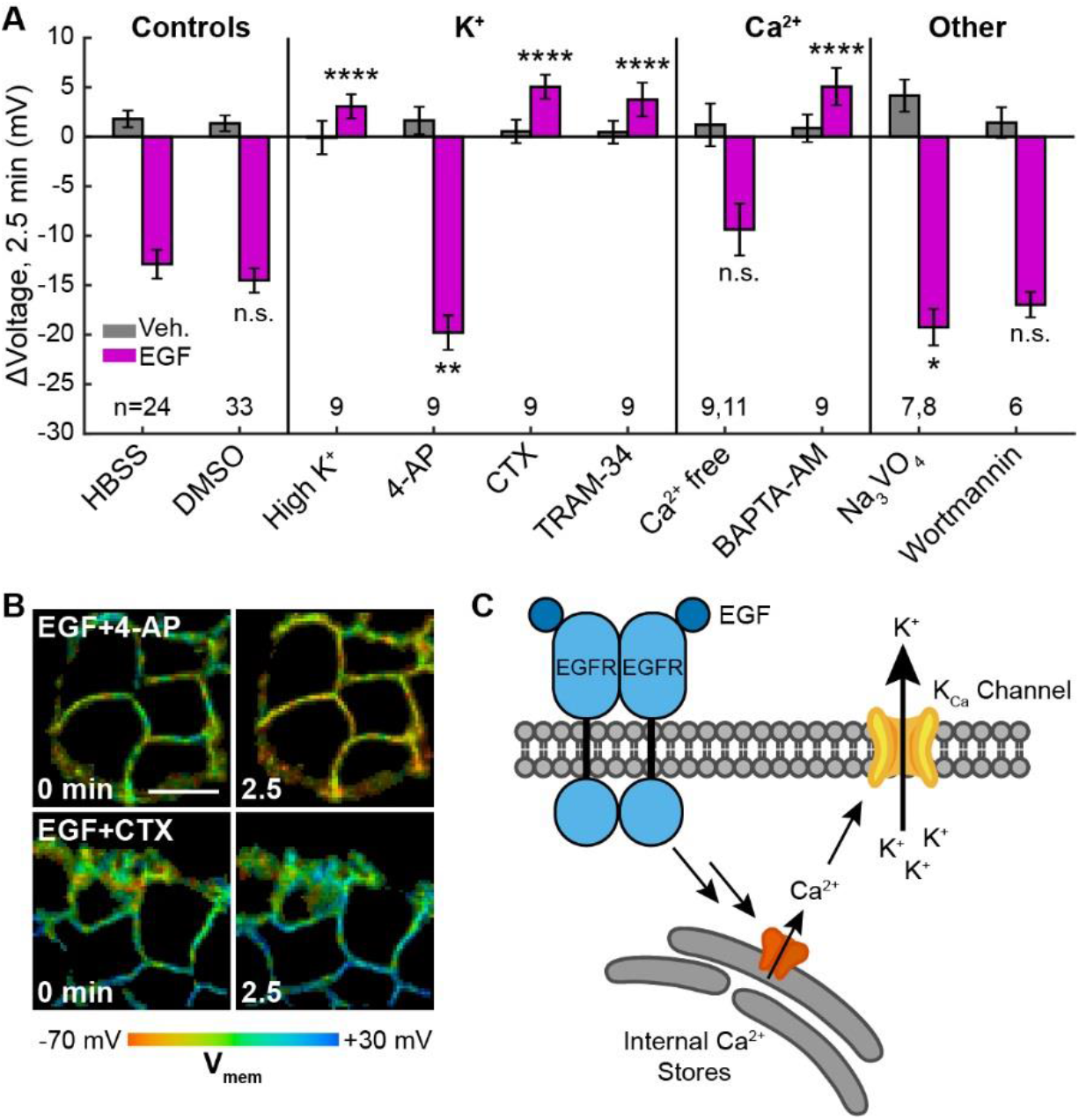
EGF-induced hyperpolarization is mediated by a Ca^2+^ activated K^+^ channel. (**A**) Comparison of the V_mem_ change 2.5 minutes after EGF addition in cells incubated in unmodified imaging buffer (HBSS) or in modified solutions. (**B**) Lifetime images of A431 cells treated with 4-AP or CTX. (**C**) Model for membrane hyperpolarization following EGFR activation. Scale bar is 20 μm. Bars are mean ± SEM. Sample sizes listed are (Veh, EGF); where only one number is given, sample size was the same for both. Asterisks reflect significant differences in EGF-stimulated V_mem_ change between the unmodified control (HBSS or DMSO) and modified solutions (n.s. p>0.05, * p<0.05, ** p<0.01, *** p<0.001, **** p<0.0001, two-tailed, unpaired, unequal variances t-test). DMSO: 0.1% DMSO, high K^+^: 120 mM K^+^, 4-AP: 5 mM 4-aminopyridine, CTX: 100 nM charybdotoxin, TRAM-34: 200 nM TRAM-34, Ca^2+^ free: 0 mM Ca^2+^ and Mg^2+^, BAPTA-AM: 10 μM bisaminophenoxyethanetetraacetic acid acetoxymethyl ester, Na3VO4: 100 μM sodium orthovanadate, wortmannin: 1 μM wortmannin.

To explore the effects of other components of the EGFR pathway on EGF-induced hyperpolarization, we perturbed intra- and extracellular Ca^2+^ concentrations during EGF stimulation. Reduction of extracellular Ca^2+^ concentration did not substantially alter the EGF response (**Fig. 5A, Fig. 5-supplement 1F**). However, sequestration of intracellular Ca^2+^ with BAPTA-AM disrupts the hyperpolarization response. BAPTA-AM treated cells show a small, 4 mV depolarization in response to EGF treatment, similar to CTX-treated cells (**Fig. 5A, Fig. 5-supplement 1G**). Perturbation of Ca^2+^ levels had little effect on the resting membrane potential (**Fig. 5-supplement 2**). Introduction of wortmannin (1 μM) to block downstream kinase activity has no effect on the membrane potential response to EGF, while orthovanadate addition (Na3VO4, 100 μM) to block phosphatase activity results in a small increase in the hyperpolarizing response (**Fig. 5A, Fig. 5-supplement 1H-I**). These results support a model for EGF-EGFR mediated hyperpolarization in which RTK activity of EGFR causes release of internal Ca^2+^ stores to in turn open KCa channels and hyperpolarize the cell (**Fig. 5C**).

## Discussion

We report the design and implementation of a new method for optically quantifying absolute membrane potential in living cells. VF-FLIM is operationally simple, requires just a single-point calibration, and is applicable across a number of cell types. VF-FLIM exhibits a 20fold improvement in voltage resolution over previous FLIM-based approaches ^20,21^, achieving sufficient resolution to make biologically relevant voltage measurements. The photoinduced electron transfer mechanism of VoltageFluors ^23^ renders superior sensitivity and consistency of the lifetime measurement; furthermore, because VoltageFluors are applied exogenously, the vast majority of the fluorescence signal is voltage-sensitive and at the membrane. Unlike small-molecule FRET-oxonol approaches to optically estimate membrane potential values ^10^, VF-FLIM presents a direct relationship between τfl and V_mem_ with a single optical reporter and avoids complex and potentially toxic multi-dye loading protocols.

Because VF-FLIM is an optical approach, it improves upon both the throughput and spatial resolution of patch clamp electrophysiology and thereby enables new lines of inquiry in biological systems. Although individual VF-FLIM measurements have more voltage-equivalent noise than modern electrophysiology, the ability to perform thousands of recordings over the course of a few hours enables a more complete documentation of the distributions of cellular V_mem_ present in a cell population. In addition to throughput, another key difference between VF-FLIM and patch-clamp electrophysiology is spatial resolution. While VF-FLIM records the V_mem_ of an optically defined region of interest (in this case a cell or cell group), electrophysiology records V_mem_ at an individual cell or part of a cell where the electrode makes contact, which may or may not reflect the V_mem_ of the entire cell or group. In principle, VF-FLIM could record subcellular differences in V_mem_ that would be difficult to dissect with electrophysiology. Looking ahead, such subcellular recordings in cells with complex morphology and processes are an exciting area for future development of VF-FLIM, in conjunction with cellular and sub-cellular strategies for targeting VF dyes ^38,39^.

We optically documented resting membrane potential distributions in cultured cells, as VF-FLIM is well suited to address questions about V_mem_ states present in these samples. The presence and significance of distinct V_mem_ states in cell populations is mostly uncharacterized due to the throughput limitations of patch-clamp electrophysiology, but some reports suggest that distinct V_mem_ states arise during the various phases of the cell cycle ^40,41^. V_mem_ histograms presented in this work appear more or less unimodal, showing no clear sign of cell cycle-related V_mem_ states (**Fig. 3A,D; Fig. 3-supplement 1A,D,G**). We considered the possibility that VF-FLIM does not detect cell-cycle-related V_mem_ states because we report average V_mem_ across cell groups in cases where cells are in contact (**Scheme S2**). This explanation is unlikely for two reasons. First, V_mem_ distributions for CHO cells appear unimodal, even though CHO cultures were mostly comprised of isolated cells under the conditions tested (**Fig. 3D-F**). Second, theoretical work suggests that dramatically different V_mem_ states in adjacent cells are unlikely, as electrical coupling often leads to equilibration of V_mem_ across the cell group ^42,43^. Although we cannot rule out the possibility of poorly separated V_mem_ populations (i.e. with a mean difference in voltage below our resolution limit), VF-FLIM both prompts and enables a re-examination of the notion that bi- or multimodal V_mem_ distributions exist in cultured cells. Furthermore, VF-FLIM represents an exciting opportunity to experimentally visualize theorized V_mem_ patterns in culture and in more complex tissues. Studies towards this end are ongoing in our laboratory.

In the present study, we use VF-FLIM to provide the first cell-resolved, direct visualization of voltage changes induced by growth factor signaling. For long term V_mem_ recordings during growth-related processes, an optical approach is more attractive than an electrode-based one. Electrophysiology becomes increasingly challenging as time scale lengthens, especially if cells migrate, and washout of the cytosol with pipette solution can change the very signals under study ^44,45^. Previous attempts to electrophysiologically record V_mem_ in EGF-stimulated A431 cells were unsuccessful due to these technical challenges.^34^ Because whole cell voltage-clamp electrophysiology was intractable, the V_mem_ response in EGF-stimulated A431 cells was addressed indirectly through model cell lines expressing EGFR exogenously ^34^, bulk measurements on trypsinized cells in suspension ^46^, or cell-attached single channel recordings ^47–49^. By stably recording V_mem_ during EGF stimulation, VF-FLIM enables direct study of V_mem_ signaling in otherwise inaccessible pathways.

In conjunction with physiological manipulations and pharmacological perturbations, we explore the molecular mechanisms underlying EGF-induced hyperpolarization. We find that signaling along the EGF-EGFR axis results in a robust hyperpolarizing current carried by K^+^ ions, passed by the Ca^2+^-activated K^+^ channel Kea3.1, and mediated by intracellular Ca^2+^ (**Fig. 5C**). We achieve a complete loss of the hyperpolarizing response to EGF by altering the K^+^ driving force (“High K^+^” **Fig. 5A, Fig. 5-supplement 1B**), blocking calcium-activated K^+^ currents directly (“CTX” and “TRAM-34”, **Fig. 5A, Fig. 5-supplement 1D,E**), or intercepting cytosolic Ca^2+^ (“BAPTA-AM”, **Fig. 5A, Fig. 5-supplement 1G**). These results, combined with transcriptomic evidence that Kea3.1 is the major Kea channel in A431 cells ^50^, indicate that Kea3.1 mediates the observed hyperpolarization. Interestingly, under some conditions where K^+^-mediated hyperpolarization is blocked (“CTX,” “high K^+^”, “BAPTA-AM”), VF-FLIM reveals a small, secondary depolarizing current not visible during normal EGF stimulation. This current likely arises from initial Ca^2+^ entry into the cell, as previously observed during EGF signaling ^51,52^. Although we have obtained direct and conclusive evidence of EGF-induced hyperpolarization in A431 cells, the interactions between this voltage change and downstream targets of EGFR remain incompletely characterized. Enhancing EGF signaling by blockade of cellular tyrosine phosphatases with orthovanadate ^53^ correspondingly increases EGF-mediated hyperpolarization (“Na3VO4” **Fig. 5A, Fig. 5-supplement 1H**), but inhibition of downstream kinase activity appears to have little effect on hyperpolarization (“wortmannin” **Fig. 5A, Fig. 5-supplement 1I**).

In the context of RTK signaling, V_mem_ may serve to modulate the driving force for external Ca^2+^ entry ^3,54^ and thereby act as a regulator of this canonical signaling ion. Alternatively, V_mem_ may play a more subtle biophysical role, such as potentiating lipid reorganization in the plasma membrane ^55^. Small changes in V_mem_ likely affect signaling pathways in ways that are currently completely unknown, but high throughput discovery of V_mem_ targets remains challenging. Combination of electrophysiology with single cell transcriptomics has begun to uncover relationships between V_mem_ and other cellular pathways in excitable cells ^56^; such approaches could be coupled to higher throughput VF-FLIM methods to explore pathways that interact with V_mem_ in non-excitable contexts.

VF-FLIM represents a novel and general approach for interrogating the roles of membrane potential in fundamental cellular physiology. Future improvements to the voltage resolution could be made by use of more sensitive indicators, which may exhibit larger changes in fluorescence lifetime ^24^ VF-FLIM can be further expanded to include the entire color palette of PeT-based voltage indicators ^57,58^, allied with targeting methods to probe absolute membrane potential in heterogeneous cellular populations ^38,39^, and coupled to high-speed imaging techniques for optical quantification of fast voltage events ^59^.

## Acknowledgments

We thank Holly Aaron and Vadim Degtyar for expert technical assistance and training in the use of FLIM, Prof. John Kuriyan and Dr. Sean Peterson for helpful discussions, and members of the Miller lab for providing VF dyes. FLIM experiments were performed at the CRL Molecular Imaging Center, supported by NSF DBI-0116016. Cell lines were from the UCB Cell Culture Facility. FCK-QuasAr2-Citrine was a gift from Adam Cohen (Addgene plasmid # 59172). JLD was supported by an NSF Graduate Research Fellowship. EWM acknowledges support from the Sloan Foundation (FG-2016-6359), March of Dimes (5-FY16-65), and the NIH (R35GM119855).

## Author Contributions

JLD performed experiments, analyzed data, and wrote the paper. AMMG performed experiments and analyzed data. EWM analyzed data and wrote the paper.

## Competing Interest Statement

EWM is listed as an inventor on a patent describing voltage-sensitive fluorophores. This patent is owned by the Regents of the University of California.

## Materials and Methods

### Materials

VoltageFluor dyes VF2.1.Cl and VF2.0.Cl were synthesized in house according to previously described syntheses ^24^. Dyes were stored either as solids at room temperature or as 1000x DMSO stocks at −20°C. Dye stock concentrations were normalized to the absorption of the dichlorofluorescein dye head via UV-Vis spectroscopy.

All salts and buffers were purchased from either Sigma-Aldrich (St. Louis, MO) or Fisher Scientific (Waltham, MA). TRAM-34, 4-aminopyridine, and charybdotoxin were purchased from Sigma-Aldrich. Gefitinib, wortmannin, sodium orthovanadate, and BAPTA-AM were purchased from Fisher Scientific. Canertinib was a gift from the Kuriyan laboratory at UC Berkeley. Gefitinib, wortmannin, canertinib, and TRAM-34 were made up as 1000x-10000x stock solutions in DMSO and stored at −20°C. Charybdotoxin was made up as a 1000x solution in water and stored at −80°C. 4-aminopyridine was made up as a 20x stock in imaging buffer (HBSS) and stored at 4°C. Recombinantly expressed epidermal growth factor was purchased from PeproTech (Rocky Hill, NJ) and aliquoted as a 1 mg/mL solution in water at −80°C.

Solid sodium orthovanadate was dissolved in water and activated before use ^60^. Briefly, orthovanadate solutions were repeatedly boiled and adjusted to pH 10 until the solution was clear and colorless. 200 mM activated orthovanadate stocks were aliquoted and stored at −20°C.

Unless otherwise noted, all imaging experiments were performed in Hank’s Balanced Salt Solution (HBSS; Gibco/Thermo Fisher Scientific). HBSS composition in mM: 137.9 NaCl, 5.3 KCl, 5.6 D-glucose, 4.2 NaHCO_3_, 1.3 CaCl_2_, 0.49 MgCl_2_, 0.44 KH2PO_4_, 0.41 MgSO4, 0.34 Na2HPO4. High K^+^ HBSS was made in-house to 285 mOsmol and pH 7.3, containing (in mM): 120 KCl, 23.3 NaCl, 5.6 D-glucose, 4.2 NaHCO_3_, 1.3 CaCl_2_, 0.49 MgCl_2_, 0.44 KH2PO_4_, 0.41 MgSO_4_, 0.34 Na_2_HPO_4_. Nominally Ca^2+^/Mg^2+^ free HBSS (Gibco) contained, in mM: 137.9 NaCl, 5.3 KCl, 5.6 D-glucose, 4.2 NaHCO_3_, 0.44 KH2PO_4_, 0.34 Na_2_HPO_4_.

## Methods

### Cell Culture

All cell lines were obtained from the UC Berkeley Cell Culture Facility and discarded after twenty passages. Cells were maintained in Dulbecco’s Modified Eagle Medium (DMEM) with 4.5 g/L D-glucose supplemented with 10% FBS (Seradigm (VWR); Radnor, PA) and 2 mM GlutaMAX (Gibco) in a 5% CO2 incubator at 37°C. Media for MCF-7 cells was supplemented with 1 mM sodium pyruvate (Life Technologies/Thermo Fisher Scientific) and 1x non-essential amino acids (Thermo Fisher Scientific). Media for CHO.K1 (referred to as CHO throughout the text) cells was supplemented with 1x non-essential amino acids. HEK293T and MDA-MB-231 were dissociated with 0.05% Trypsin-EDTA with phenol red (Thermo Fisher Scientific) at 37°C, whereas A431, CHO, and MCF-7 cells were dissociated with 0.25% Trypsin-EDTA with phenol red at 37°C. To avoid potential toxicity of residual trypsin, all cells except for HEK293T were spun down at 250xg or 500xg for 5 minutes and re-suspended in fresh complete media during passaging.

For use in imaging experiments, cells were plated onto 25 mm diameter poly-D-lysine coated #1.5 glass coverslips (Electron Microscopy Sciences) in 6 well tissue culture plates (Corning; Corning, NY). To maximize cell attachment, coverslips were treated before use with 1-2 M HCl for 2-5 hours and washed overnight three times with 100% ethanol and three times with deionized water. Coverslips were sterilized by heating to 150°C for 2-3 hours. Before use, coverslips were incubated with poly-D-lysine (Sigma-Aldrich, made as a 0.1 mg/mL solution in phosphate-buffered saline with 10 mM Na3BO4) for 1-10 hours at 37°C and then washed twice with water and twice with Dulbecco’s phosphate buffered saline (dPBS, Gibco).

A431, CHO, HEK293T, and MCF-7 were seeded onto glass coverslips 16-24 hours before microscopy experiments. MDA-MB-231 cells were seeded 48 hours before use because it facilitated formation of gigaseals during whole-cell voltage clamp electrophysiology. Cell densities used for optical resting membrane potential recordings (in 10^3^ cells per cm^2^) were: A431 42; CHO 42; HEK293T 42; MCF-7 63; MDA-MB-231 42. To ensure the presence of single cells for whole-cell voltage clamp electrophysiology, fast-growing cells were plated more sparsely (approximately 20% confluence) for electrophysiology experiments. Cell densities used for electrophysiology (in 10^3^ cells per cm^2^) were: A431 36-52; CHO 21; HEK293T 21; MCF-7 63; MDA-MB-231 42. To reduce their rapid growth rate, HEK293T cells were seeded onto glass coverslips in reduced glucose (1 g/L) DMEM with 10% FBS, 2 mM GlutaMAX, and 1 mM sodium pyruvate for electrophysiology experiments.

### Cellular Loading of VoltageFluor Dyes

Cells were loaded with 1x VoltageFluor in HBSS for 20 minutes in a 37°C incubator with 5% CO2. For most experiments, 100 nM VoltageFluor was used. Serum-starved A431 cells were loaded with 50 nM VoltageFluor. After VF loading, cells were washed once with HBSS and then placed in fresh HBSS for imaging. All imaging experiments were conducted at room temperature under ambient atmosphere. Cells were used immediately after loading the VF dye, and no cells were kept for longer than an hour at room temperature.

### Whole-Cell Patch-Clamp Electrophysiology

Pipettes were pulled from borosilicate glass with filament (Sutter Instruments, Novato, CA) with resistances ranging from 4 to 7 MΩ with a P97 pipette puller (Sutter Instruments). Internal solution composition, in mM (pH 7.25, 285 mOsmol/L): 125 potassium gluconate, 10 KCl, 5 NaCl, 1 EGTA, 10 HEPES, 2 ATP sodium salt, 0.3 GTP sodium salt. EGTA (tetraacid form) was prepared as a stock solution in either 1 M KOH or 10 M NaOH before addition to the internal solution. Pipettes were positioned with an MP-225 micromanipulator (Sutter Instruments). A liquid junction potential of − 14 mV was determined by the Liquid Junction Potential Calculator in the pClamp software package ^61^ (Molecular Devices, San Jose, CA), and all voltage step protocols were corrected for this offset.

Electrophysiology recordings were made with an Axopatch 200B amplifier and digitized with a Digidata 1440A (Molecular Devices). Signals were filtered with a 5 kHz low-pass Bessel filter. Correction for pipette capacitance was performed in the cell attached configuration. Voltage-lifetime calibrations were performed in V-clamp mode, with the cell held at the potential of interest for 15 or 30 seconds while lifetime was recorded. Potentials were applied in random order, and membrane test was conducted between each step to verify the quality of the patch. For single cell patching, recordings were only included if they maintained a 30:1 ratio of membrane resistance (Rm) to access resistance (Ra) and an Ra value below 30 MΩ throughout the recording. Due to the reduced health of HEK293T cells transfected with CAESR, recordings were used as long as they maintained a 10:1 Rm:Ra ratio, although most recordings were better than 30:1 Rm:Ra. Only recordings stable for at least 4 voltage steps (roughly 2 minutes) were included in the dataset.

For electrophysiology involving small groups of cells (**Fig. 2-supplement 4**), complete voltage clamp across the entire cell group was not possible. Recordings were used as long as R_a_ remained below 30 MΩ for at least three voltage steps. Most recordings also retained Rm:Ra ratios greater than 20:1.

### Epidermal Growth Factor Treatment

A431 cells were serum starved prior to epidermal growth factor studies. Two days before the experiment, cells were trypsizined and suspended in complete media with 10% FBS. Cells were then spun down for 5 minutes at 500xg and re-suspended in reduced serum DMEM (2% FBS, 2 mM GlutaMAX, 4.5 g/L glucose). Cells were seeded onto 25 mm coverslips in 6 well plates at a density of 84 × 10^3^ cells per cm^2^. 4-5.5 hours before the experiment, the media was exchanged for serum-free DMEM (0% FBS, 2 mM GlutaMAX, 4.5 g/L glucose).

After 4-5.5 hours in serum-free media, cells were loaded with 50 nM VF dye as described above. In pharmacology experiments, the drug or vehicle was also added to the VF dye loading solution. All subsequent wash and imaging solutions also contained the drug or vehicle. For changes to buffer ionic composition, VoltageFluor dyes were loaded in unmodified HBSS to avoid toxicity from prolonged incubation with high K^+^ or without Ca^2+^. Immediately prior to use, cells were washed in the modified HBSS (120 mM K^+^ or 0 mM Ca^2+^) and recordings were made in the modified HBSS.

For analysis of short-term responses to EGF (3 minute time series), VF lifetime was recorded in 6 sequential 30 second exposures. Immediately after the conclusion of the first frame (30-35 seconds into the recording), EGF or vehicle (imaging buffer only) was added to the indicated final concentration from a 2x solution in HBSS imaging buffer. For analysis of longterm responses to EGF (15 minute time series), EGF addition occurred in the same way, but a gap of 150 seconds (without laser illumination) was allotted between each 30 second lifetime recording. Times given throughout the text correspond to the start of an exposure. Voltage changes at 2.5 minutes were calculated from the difference between an initial image (taken before imaging buffer vehicle or EGF addition) and a final image (a 30 second exposure starting 2.5 minutes into the time series).

### Transfection and Imaging of CAESR in HEK293T

The CAESR plasmid was obtained as an agar stab (FCK-Quasar2-Citrine, Addgene #59172), cultured overnight in LB with 100 μg/mL ampicillin, and isolated via a spin miniprep kit (Qiagen). HEK293T cells were plated at a density of 42,000 cells per cm^2^ directly onto a 6 well tissue culture plate and incubated at 37°C in a humidified incubator for 24 hours prior to transfection. Transfections were performed with Lipofectamine 3000 according to the manufacturer’s protocol (Thermo Fisher Scientific). Cells were allowed to grow an additional 24 hours after transfection before they were plated onto glass coverslips for microscopy experiments (as described above for electrophysiology of untransfected HEK293T cells).

### Determination of EC50 for EGF in A431 Cells

Average voltage changes 2.5 minutes after addition of EGF to serum deprived A431 cells were determined at different EGF concentrations, and these means were fit to a four parameter logistic function in MATLAB (MathWorks, Natick, MA).

### Goldman-Hodgkin-Katz Estimation of V_mem_ in Different Imaging Buffers

If intracellular and extracellular concentrations, as well as relative permeabilities, of all ionic species are known, the Goldman-Hodgkin-Katz (GHK) equation (eqn. 1) can be used to calculate the resting membrane potential of a cell ^32^. In practice, the intracellular ion concentrations [X]in and relative permeabilities Px are difficult to determine. To obtain a range of reasonable V_mem_ values in systems where these concentrations and relative permeabilities are not known, we calculated possible V_mem_ using the “standard” parameters derived from the work of Hodgkin and Katz ^32^, as well as a value above and a value below each “standard” point. The values evaluated were the following: P_k_ 1; P_Na_ 0.01, 0.05, 0.2; P_cl_ 0.2, 0.45, 0.9; [K^+^]in 90, 150, 200 mM; [Na^+^]in 5, 15, 50 mM; [C1_−_]_in_ 2, 10, 35 mM. Extracellular ion concentrations [X]_out_ were known (see **Materials**). In eqn. 1, R is the universal gas constant, T is the temperature (293 K for this experiment), and F is Faraday’s constant.

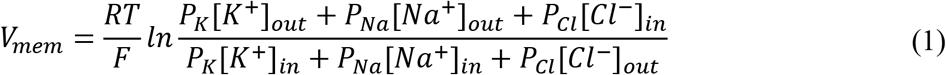

### Fluorescence Lifetime Data Acquisition

Fluorescence lifetime imaging was conducted on a LSM 510 inverted scanning confocal microscope (Carl Zeiss AG, Oberkochen, Germany) equipped with an SPC-150 or SPC-150N single photon counting card (Becker & Hickl GmbH, Berlin, Germany) (**Scheme S1**). 80 MHz pulsed excitation was supplied by a Ti:Sapphire laser (MaiTai HP; SpectraPhysics, Santa Clara, CA) tuned to 958 nm and frequency-doubled to 479 nm. The laser was cooled by a recirculating water chiller (Neslab KMC100). Excitation light was directed into the microscope with a series of silver mirrors (Thorlabs, Newton, NJ or Newport Corporation, Irvine, CA).

Excitation light power at the sample was controlled with a neutral density (ND) wheel and a polarizer (P) followed by a polarizing beamsplitter (BS). Light was titrated such that VoltageFluor lifetime did not drift during the experiment, no phototoxicity was visible, and photon pile-up was not visible on the detector. For recordings at high VoltageFluor concentrations (**Fig. 1-supplement 3, Fig. 2-supplement 5**), reduced power was used to avoid saturating the detector. For optical voltage determinations using 50 or 100 nM VoltageFluor, typical average power at the sample was 5 μW.

Fluorescence emission was collected through a 40x oil immersion objective (Zeiss) coated with immersion oil (Immersol 518F, Zeiss). Emitted photons were detected with a hybrid detector, HPM-100-40 (Becker & Hickl), based on a Hamamatsu R10467 GaAsP hybrid photomultiplier tube. Detector dark counts were kept below 1000 per second during acquisition. Emission light was collected through a 550/49 bandpass filter (Semrock, Rochester, NY) after passing through a 488 LP dichroic mirror (Zeiss). The reference photons for determination of photon arrival times were detected with a PHD-400-N high speed photodiode (Becker & Hickl). Data were acquired with 256 time bins in the analog-to-digital-converter and either 64×64 or 256×256 pixels of spatial resolution (see discussion of pixel size below).

Routine evaluation of the proper functioning of the lifetime recording setup was performed by measurement of three standards (**Fig. 1-supplement 2**): 2 μM fluorescein in 0.1 N NaOH, 1 mg/mL erythrosin B in water (pH 7), and the instrument response function (IRF). The IRF was determined from a solution of 500 μM fluorescein and 12.2 M sodium iodide in 0.1 N NaOH. Because of the high concentration of iodide quencher, the IRF solution has a lifetime shorter than the detector response time, allowing approximation of the instrument response function under identical excitation and emission conditions as data acquisition ^62^.

**Scheme S1.**
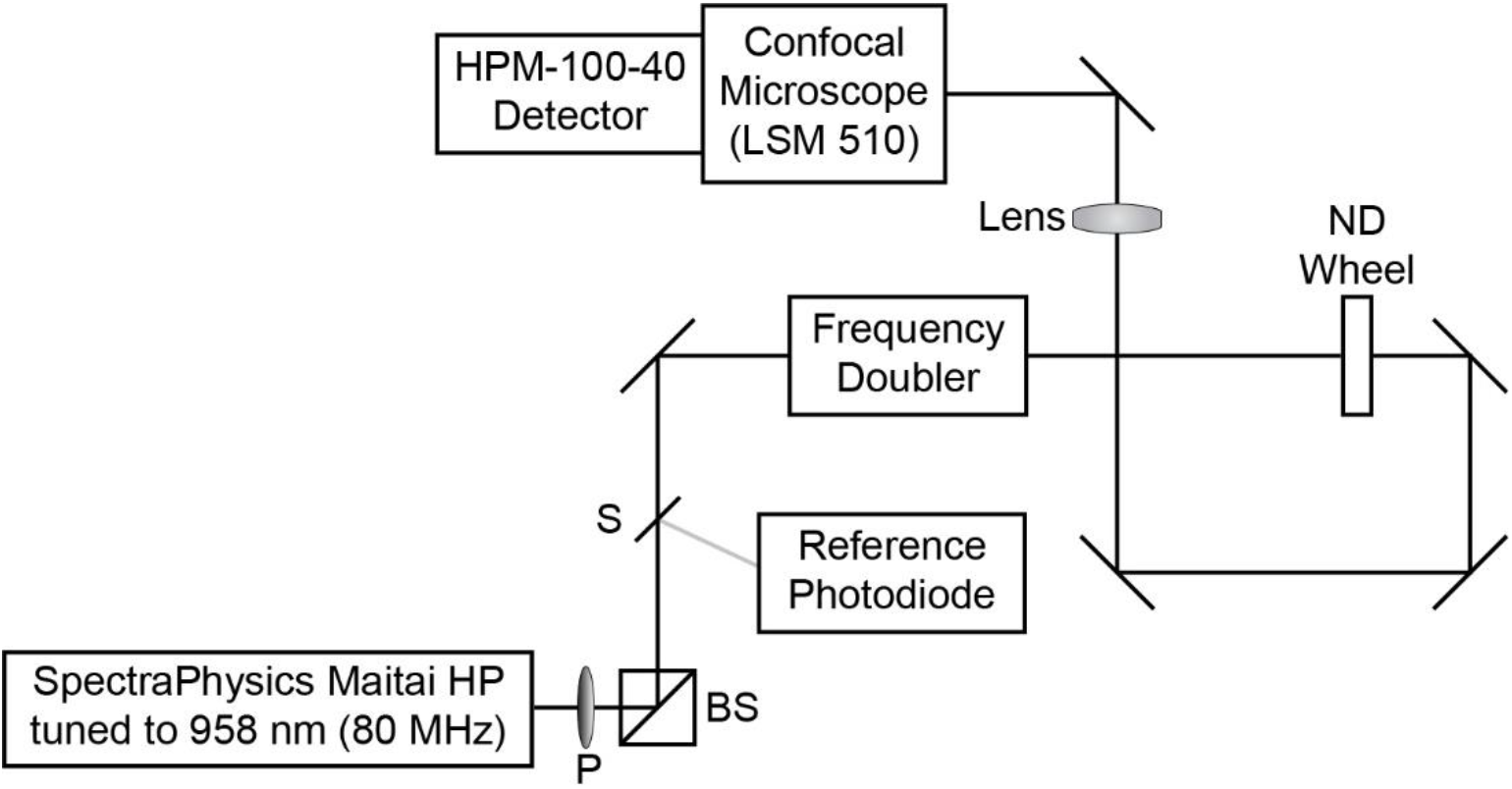
Optical diagram for time correlated single photon counting microscope. Excitation light was supplied by a Ti:Sapphire laser tuned to 958 nm. A small amount of light was redirected by a beam sampler (S) to a reference photodiode. The remaining light was passed through a frequency doubler to obtain 479 nm excitation light, which entered the LSM510 confocal microscope. A polarizer (P) followed by a polarizing beamsplitter (BS), as well as a neutral density (ND) wheel, allowed control of the amount of light passed to the sample.

### Fluorescence Lifetime Data Processing and Conversion to Voltage

**Scheme S2.**
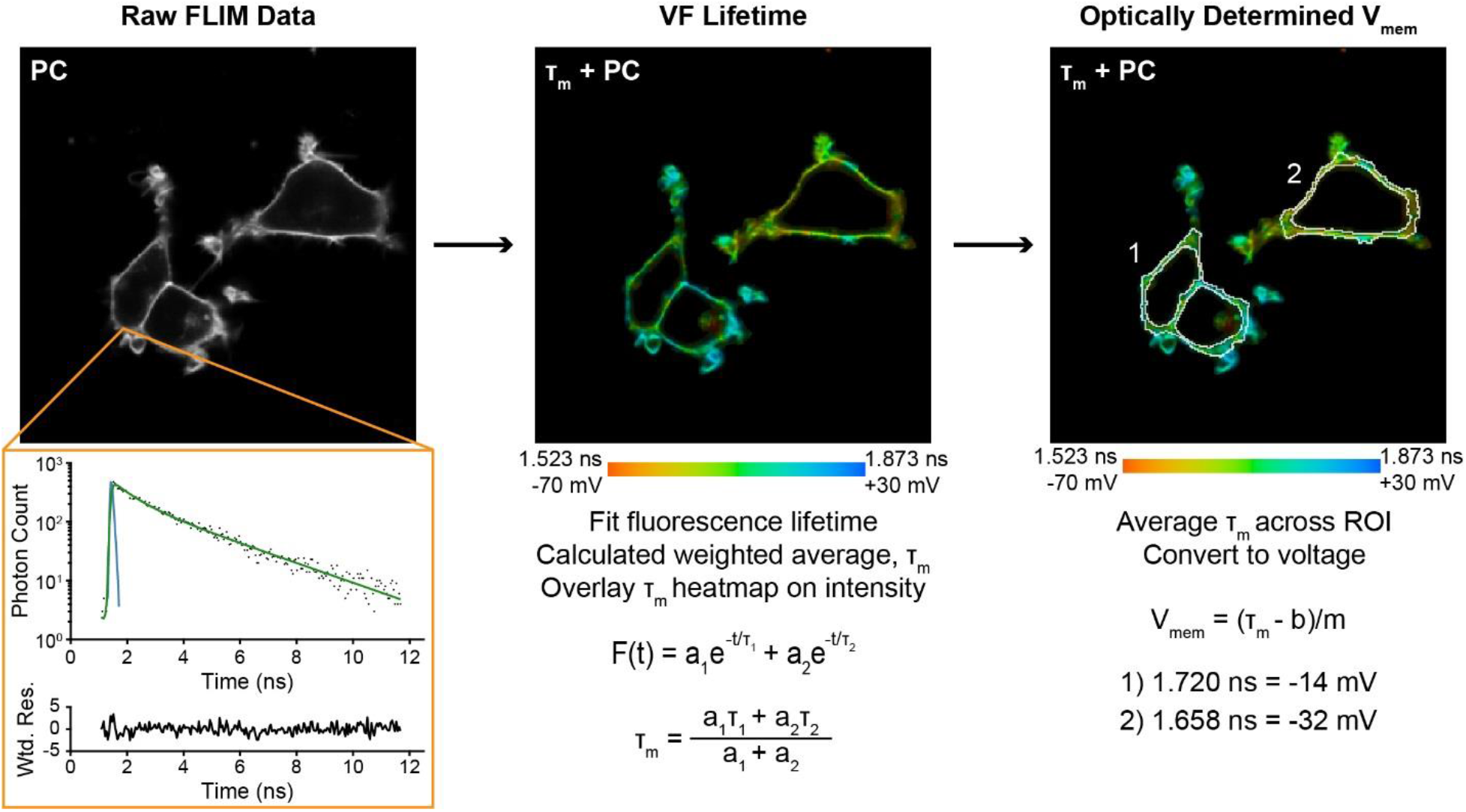
Overview of data processing to obtain membrane potential recordings from fluorescence lifetime. Time-correlated photon data (black dots, first panel) collected at each pixel were fit to an exponential decay model (green) with iterative reconvolution of the instrument response function (IRF, blue). The two components of the fluorescence lifetimes were converted to a weighted average (middle panel). Cell membranes (white outlines) were identified, and τm was averaged within each of these regions of interest (ROIs, right panel). These lifetimes were then converted to voltage via a previously determined lifetime-V_mem_ standard curve with slope m and y-intercept b. Additional details of this process are provided in the text below. Wtd. Res.: weighted residuals of the fit, τm: weighted average fluorescence lifetime, PC: photon count. τm + PC represents an overlay of the lifetime data (color heat map) onto the photon count image.

### IRF Deconvolution

Signal from photons detected in a TCSPC apparatus are convolved with the instrument response (IRF). IRFs can be approximated by the SPCImage fitting software, but consistency of lifetime fits on VF-FLIM datasets was improved by using a measured IRF. Measured IRFs were incorporated by the iterative reconvolution method using SPCImage analysis software ^63^.

### VoltageFluor Lifetime Fitting Model

All VoltageFluor lifetime data were fit using SPCImage (Becker & Hickl), which solves the nonlinear least squares problem using the Levenberg-Marquadt algorithm. VF2.1.Cl lifetime data were fit to a sum of two exponential decay components (eqn. 2). Attempts to fit the VF2.1.Cl data with a single exponential decay (eqn. 3) were unsatisfactory.

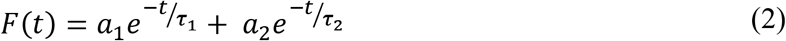

The fluorescence lifetime of VF2.0.Cl was adequately described by a single exponential decay for almost all data (eqn. 3). A second exponential component was necessary to fit data at VF2.0.Cl concentrations above 500 nM, likely attributable to the concentration-dependent decrease in lifetime that was observed high VF concentrations.

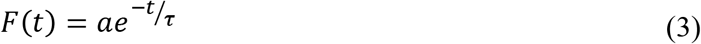

For all data fit with the two component model, the weighted average of the two lifetimes, τ_m_ (eqn. 4), was used in subsequent analysis.

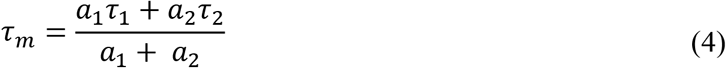

All lifetime images are represented as an overlay of photon count (pixel intensity) and weighted average lifetime (pixel color) throughout the text (τ_m_ + PC, **Scheme S2**). Pixels with insufficient signal to fit a fluorescence decay are shown in black. The photon counts, as well as the lifetimes, in image sequences on the same set of cells are scaled across the same range.

### Additional Fit Parameters for VoltageFluor Lifetimes

Pixels with photon counts below 300 (VF2.1.Cl) or 150 (VF2.0.Cl) photons at the peak of the decay (time bin with the most signal) were omitted from analysis to ensure reproducible fits. Because the lifetime of VFs does not fully decay to baseline in a single 12.5 ns laser cycle, the incomplete multiexponentials fitting option was used, allowing the model to attribute some signal early in the decay to the previous laser cycle. Out of 256 time bins from the analog-to-digital converter (ADC), only data from time bins 23 to 240 were used in the final fit. The offset parameter (detector dark counts per ADC time bin per pixel) was set to zero. The number of iterations for the fit in SPCImage was increased to 20 to obtain converged fits. Shift between the IRF and the decay trace was fixed to 0.5 (in units of ADC time bins), which consistently gave lifetimes of standards erythrosin B (1 mg/mL in H2O) ^64^ and fluorescein (2 μM in 0.1 N NaOH, H2O) ^27^ closest to reported values (**Fig. 1 – supplement 2**).

### Effective Pixel Size

To obtain sufficient photons but keep excitation light power minimal, binning between neighboring pixels was employed during fitting. This procedure effectively takes the lifetime as a spatial moving average across the image by including adjacent pixels in the decay for a given pixel.

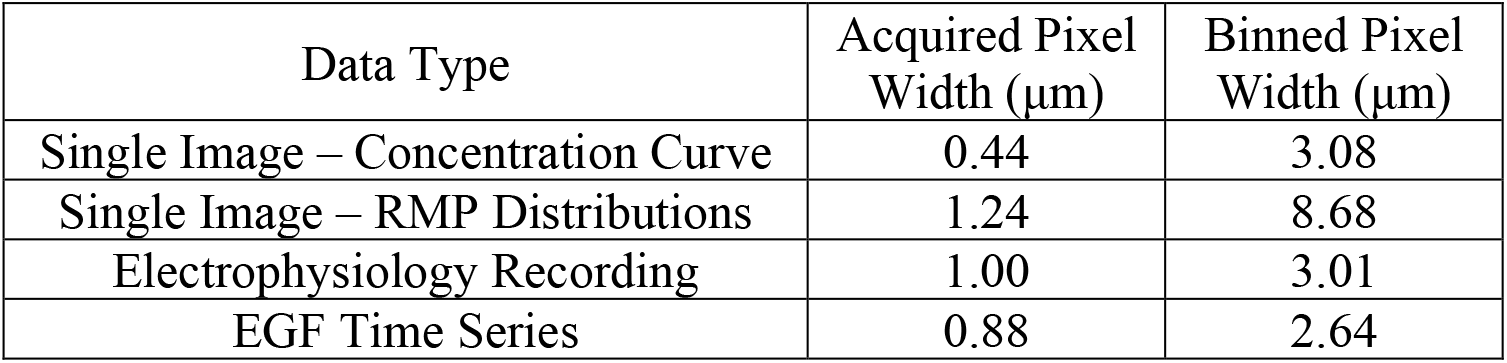

| Data Type | Acquired Pixel Width ($\mu\text{m}$) | Binned Pixel Width ($\mu\text{m}$) |
| --- | --- | --- |
| Single Image – Concentration Curve | 0.44 | 3.08 |
| Single Image – RMP Distributions | 1.24 | 8.68 |
| Electrophysiology Recording | 1.00 | 3.01 |
| EGF Time Series | 0.88 | 2.64 |

**Pixel Sizes.** For each recording type, the width of each pixel at acquisition is reported, as well as the width of the area included in the binned lifetime signal during fitting. All pixels are square.

### Determination of Regions of Interest

Images were divided into cell groups, with each cell group as a single region of interest (ROI). ROIs were determined from photon count images, either manually from the cell morphology in ImageJ or automatically by sharpening and then thresholding the signal intensity with custom MATLAB code. Regions of images that were partially out of the optical section or contained punctate debris were omitted. Sample ROIs are shown in **Scheme S2**.

For cells that adjoin other cells, attribution of a membrane region to one cell versus the other is not possible. As such, we chose to interpret each cell group as an independent sample (‘n’) instead of extracting V_mem_ values for individual cells. Adjacent cells in a group are electrically coupled to varying degrees, and their resting membrane potentials are therefore not independent ^31^. While this approach did not fully utilize the spatial resolution of VF-FLIM, it prevented overestimation of biological sample size for the effect in question.

### Conversion of Lifetime to Transmembrane Potential

The mean τm across all pixels in an ROI was used as the lifetime for that ROI. Lifetime values were mapped to transmembrane potential via the lifetime-V_mem_ standard curves determined with whole-cell voltage-clamp electrophysiology. For electrophysiology measurements, the relationship between the weighted average lifetime (eqn. 4) and membrane potential for each patched cell was determined by linear regression, yielding a sensitivity (*m*, ps/mV) and a 0 mV lifetime (*b*, ps) for each cell (eqn. 5). The average sensitivity and 0 mV point across all cells of a given type were used to convert subsequent lifetime measurements (τ) to V_mem_ (**Figure 2-supplement 3**, eqn. 6). For quantifying changes in voltage (ΔV_mem_) from changes in lifetime (Δτ), only the average sensitivity is necessary (eqn. 7).

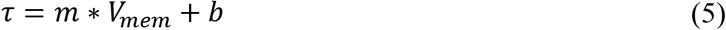

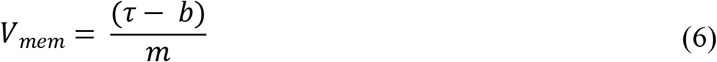

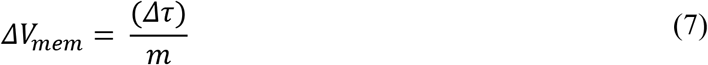

Where standard error of the mean of a voltage determination (δV_mem_) is given, error was propagated to include the standard errors of the slope (δ*m*) and y-intercept (δ*b*) of the voltage calibration, as well as the standard error of the lifetime measurements (δτ) in the condition of interest (eqn. 8). For error in a voltage change (δΔV_mem_), only error in the calibration slope was included in the propagated error (eqn. 9). Where standard deviation of VF-FLIM derived V_mem_ values is shown, a similar error propagation procedure was applied, using the standard deviation of the average sensitivity and 0 mV lifetime for that cell line.

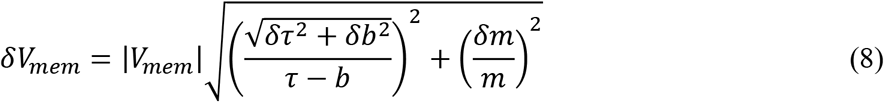

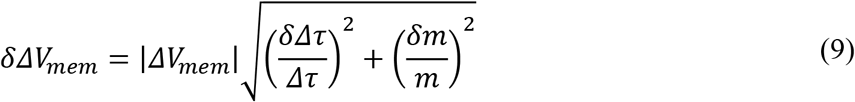

### Resolution of VF-FLIM Voltage Determination

The intrinsic nature of fluorescence lifetime introduces a point of reference into the voltage measurement, from which a single lifetime image can be interpreted as resting membrane potential. The reproducibility of this reference point (reported here as the 0 mV lifetime) over time and across cells determines the accuracy of optical V_mem_ measurements. Because the sensitivities exhibited little variability within each cell type, the slope parameter contributes very little to the overall error.

The amount of voltage-independent noise in VF-FLIM can be estimated from lifetime-V_mem_ calibration data. We report resolution as the root-mean-square deviation (RMSD) of the optically calculated voltage (V_FLIM_) from the voltage set by whole-cell voltage clamp (Vephys). The RMSD of n measurements (eqn. 10) can be determined from the variance σ^2^ (eqn. 11) and the bias (eqn. 12) of the estimator (in this case, VF-FLIM) relative to the “true” value (in this case, electrophysiology).

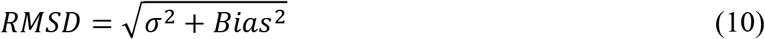

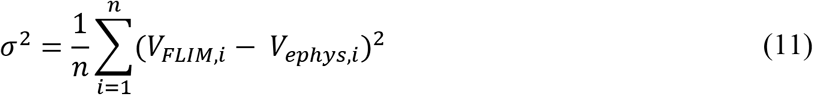

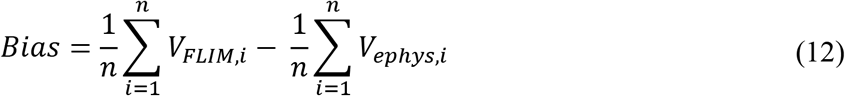

The voltage-independent variations in lifetime are much larger between cells than within a cell. Therefore, the error in tracking the magnitude of voltage changes on an individual cell (“intracell” comparisons) is much lower than the error in making a comparison of absolute V_mem_ between two cells (“inter-cell” comparisons). We can therefore determine an “intra-cell” RMSD and an “inter-cell” RMSD to reflect the voltage resolution of these two types of measurements. To calculate “intra-cell” error, we look at the RMSD between Vephys and V_FLIM_ using the τfl-V_mem_ relationship *for that specific cell*. Phrased another way, we are looking at the amount of error that would be expected in estimating a new V_mem_ for a cell based on a previous, optically-determined potential at that cell (i.e. changes in voltage). By averaging these “intra cell” RMSD values across all cells of a given type, we estimate the single-trial resolution for quantifying voltage changes is at or below 5 mV (**Fig. 2-supplement 3**).

The error in the absolute membrane potential determination (“inter-cell”) is calculated here as the RMSD between the y-intercept (0 mV lifetime) of all of the individual cells’ lifetime-voltage relationships and the 0 mV value for the averaged calibration *for all cells of a given type*. This metric addresses how well the lifetime-V_mem_ relationship for a given cell type is likely to represent an individual cell’s lifetime-V_mem_ relationship. This “inter cell” RMSD ranged from 11 to 24 mV for the tested cell lines (**Fig. 2-supplement 3**). Because of the improved throughput of VF-FLIM, much smaller errors for a population value of V_mem_ can be obtained by and averaging V_mem_ recordings from multiple cells.

This method of calculating error assumes that the electrophysiology measurement is perfectly accurate and precise. Realistically, it is likely that some of the variation seen is due to the quality of the voltage clamp. As a result, these RMSD values provide a conservative upper bound for the voltage errors in VF-FLIM.

### Analysis of CAESR Lifetimes

For sample images of CAESR in HEK293T (**Fig. 1-supplement 5**), fluorescence decays were fit using SPCImage to a biexponential decay model as described for VF2.1.Cl above, using a peak photon threshold of 150 and a bin of 2 (binned pixel width of 5 μm). To better match the studies by Cohen and co-workers ^21^, which isolated the membrane fluorescence from cytosolic fluorescence by directing the laser path, the lifetime-voltage relationships were not determined with these square-binned images. Instead, membranes were manually identified, and the fluorescence decays from all membrane pixels were summed together before fitting once per cell. (This is in contrast to the processing of VoltageFluor data, where the superior signal to noise and localization enables fitting and analysis of the lifetime on a pixel by pixel basis). This “one fit per membrane” analysis of CAESR was performed in custom MATLAB code implementing a Nelder-Meade algorithm, in which CAESR data were fit to a biexponential model with the offset fixed to 0 and the color shift as a free parameter.

## Supplementary Information

Supplementary Information for:

Optical determination of absolute membrane potential

Julia R. Lazzari-Dean, Anneliese M. M. Gest, Evan W. Miller

### Figure 1 Supplements

**Fig. 1, S1.**
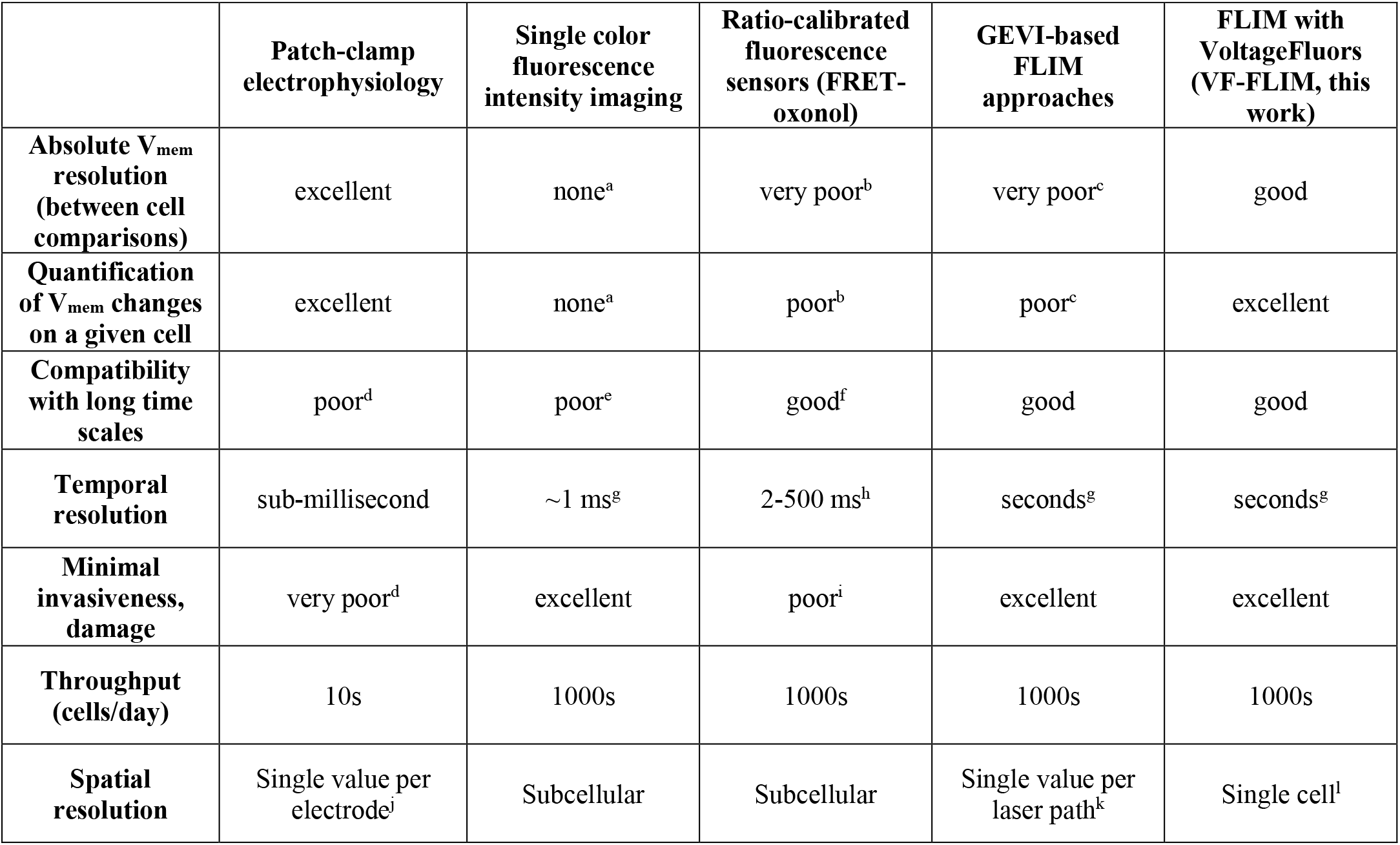
Comparison of available approaches for measuring membrane potential in cells. ^a^Measurements vary too much to be converted to absolute voltage or interpreted across populations of cells. This variability is attributable to numerous confounding factors, including dye loading, photobleaching, and sample movement ^9^. ^b^While in principle less variable than a single-color fluorescence intensity measurement, in practice, the signal depends strongly on the loading of two independent lipophilic indicators ^12,65^, which can vary substantially. ^c^Poor protein trafficking leads to large amounts of non-voltage-sensitive signal, which contaminates the FLIM recording. Voltage-equivalent resolution on a single cell (intra-cell) was 30 mV; comparisons between cells (inter-cell) show voltage-equivalent resolution of 400 mV (**Fig. 1-supplement 5, Methods**). ^d^Patch-clamp electrophysiology requires physical contact with the cell of interest, which causes damage to the cell and, in whole cell configurations, washout of intracellular factors. Slight movement of the cell or sample generally result in loss of the patch. ^e^Movement of the cell and photobleaching of the dye both cause large changes to the signal over seconds to minutes. ^f^Ratio-calibrated imaging approaches use a second signal (usually another color of fluorescence) to correct for the cell movement that contaminates single-color intensity signals over time. If the rate of photobleaching is the same for both components, photobleaching artifacts can also be avoided. ^g^Limited by photon count rates. Large numbers of photons per pixel must be collected to fit TCSPC FLIM data, leading to slower acquisition speeds. ^h^Limited by probe movement in the membrane, which depends mostly on lipophilicity ^11^. Toxicity from capacitive load of the sensor ^11^. ^j^The spatial resolution of electrophysiology is compromised by space clamp error, preventing interpretation of V_mem_ in regions far from the electrode (e.g. many neuronal processes) ^29,30^. ^k^As demonstrated by Cohen and co-workers ^20,21^; in our hands with CAESR, we also experienced significant improvements in voltage resolution by fitting a single curve per FLIM image instead of processing the images pixel-wise (see **Methods**) ^l^In this work, we calibrated VF-FLIM for V_mem_ measurements with single cell resolution. In principle, subcellular spatial resolution could be achieved with the VF-FLIM technique.

**Fig. 1, S2.**
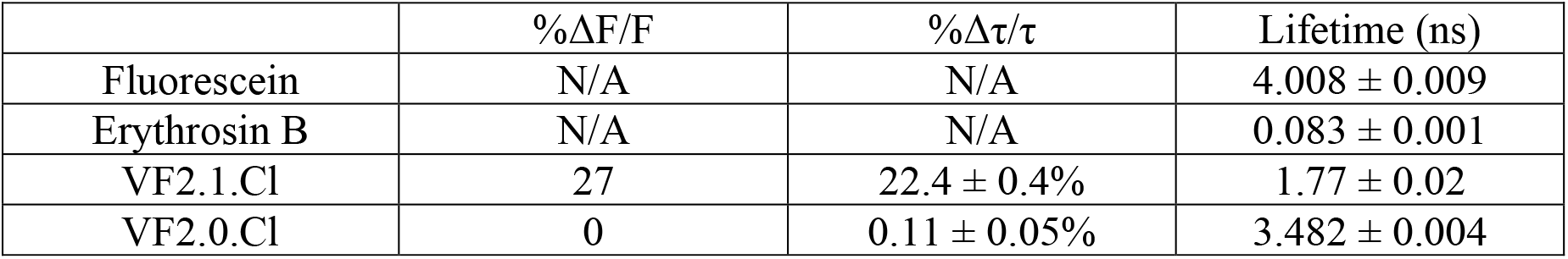
Properties of lifetime standards and VoltageFluor dyes. Fluorescein and erythrosin B standards were measured in drops of solution placed on a coverslip. For VF dyes, voltage sensitivities from intensity-based fluorescence imaging in HEK293T cells (%ΔF/F, percent change in fluorescence intensity for a voltage step from −60 mV to +40 mV) are from previously published work ^24^ Lifetime data were obtained from voltage-clamp electrophysiology of HEK293T cells loaded with 100 nM VF. Lifetime listed here is the average 0 mV lifetime from the electrophysiology calibration. % Δτ/τ is the percent change in lifetime corresponding to a 100 mV step from −60 mV to +40 mV. Lifetime sample sizes: fluorescein 25, erythrosin B 25, VF2.1.Cl 17, VF2.0.Cl 17. For lifetime standards, each measurement was taken on a separate day. VF2.1.Cl data in HEK293T is duplicated in **Figure 2 – supplement 3**. Values are tabulated as mean ± SEM.

**Fig. 1, S3.**
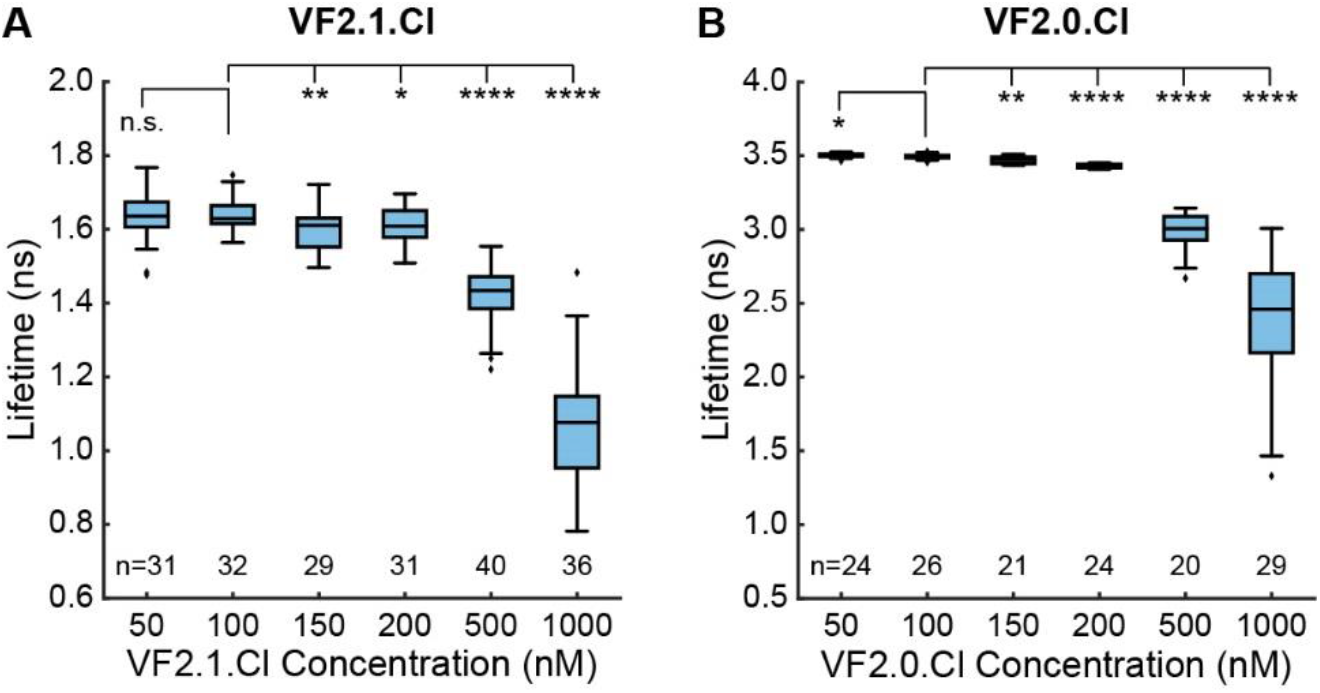
Concentration dependence of VoltageFluor lifetimes in HEK293T cells. Changes in lifetime arising from addition of a range of concentrations of (**A**) VF2.1.Cl or (**B**) voltage-insensitive control VF2.0.Cl in HEK293T cells. Biexponential fit models were used for all VF2.1.Cl concentrations and 1 μM VF2.0.Cl; a monoexponential model was used for all other VF2.0.Cl concentrations. Box plots represent the interquartile range, with whiskers and outliers determined with the Tukey method. Sample sizes indicate number of cell groups. Data were obtained over 2 to 4 different days from a total of 3 or 4 coverslips at each concentration. Asterisks indicate significant differences between the indicated concentration and the VF concentration used for electrophysiology experiments (n.s. p > 0.05, * p < 0.05, ** p < 0.01, *** p < 0.001, **** p < 0.0001, two-sided, unpaired, unequal variances t-test).

**Fig. 1, S4.**
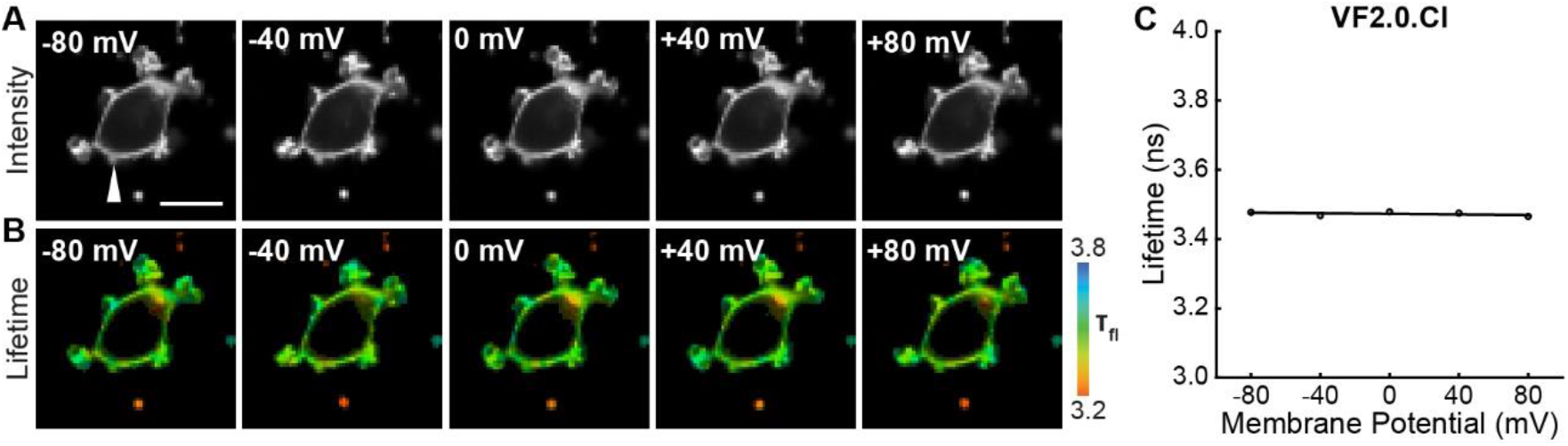
VF2.0.Cl lifetime does not depend on membrane potential. (**A**) Photon count and (**B**) lifetime images of a single HEK293T cell loaded with 100 nM VF2.0.Cl, with the membrane potential held at the indicated value via whole-cell voltage clamp electrophysiology. White arrow indicates patch pipette. Scale bar is 20 μm. (**C**) Quantification of images shown in (B) for this individual cell. Black line is the line of best fit.

**Fig. 1, S5.**
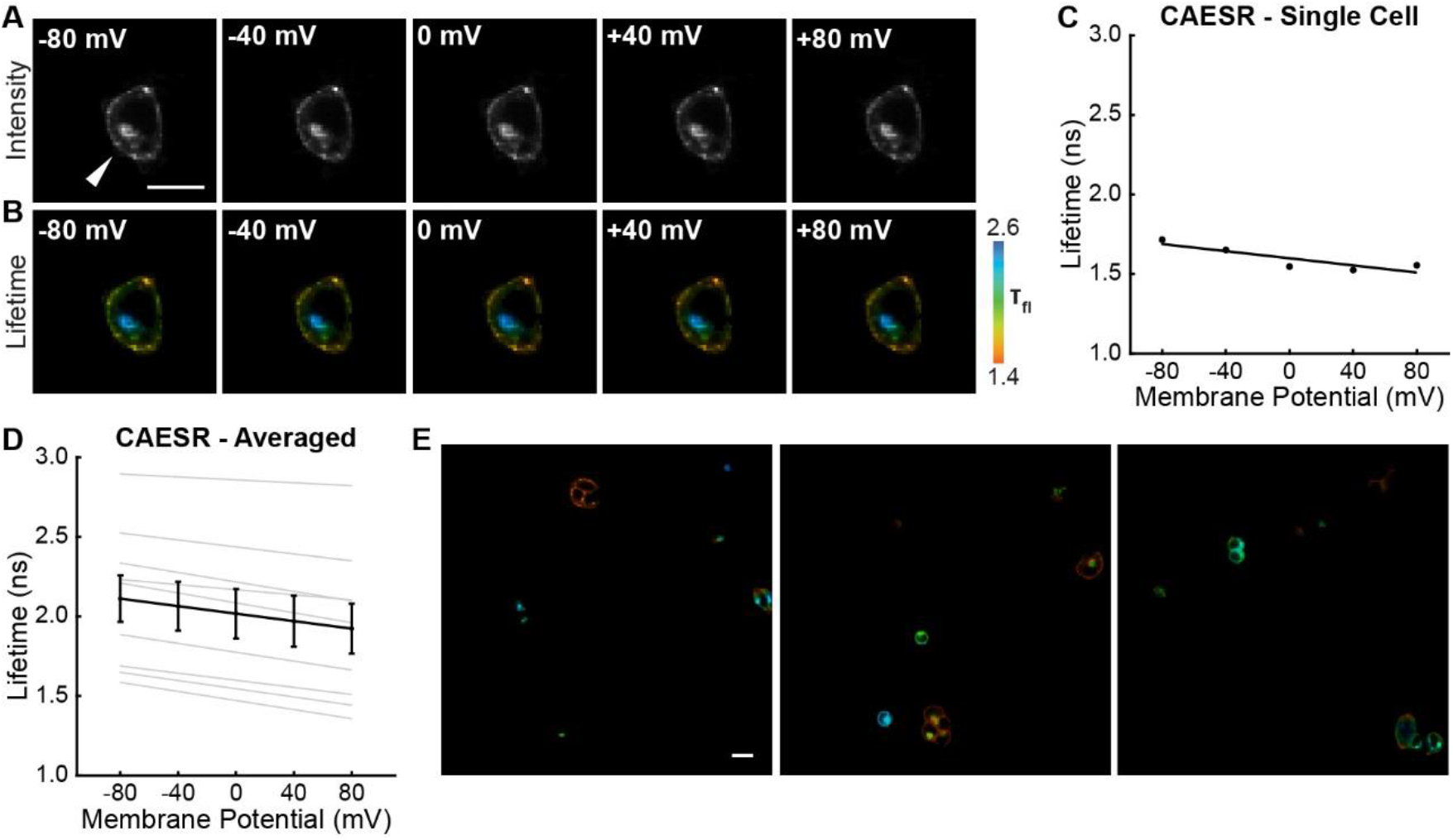
The GEVI CAESR shows variable lifetime-voltage relationships. (**A**) Photon count and (**B**) lifetime images of a HEK293T cell expressing CAESR and held at the indicated V_mem_ with voltage-clamp electrophysiology. White arrow indicates voltage-clamped cell. (**C**) Lifetime-V_mem_ relationship from the cell in (B), based on a single fit from combined fluorescence decays of all pixels in the cell membrane at each potential (see **Methods**). Points indicate recordings at a given potential; solid line is line of best fit. (**D**) Evaluation of VF2.1.Cl lifetime-voltage relationships in many individual CAESR-expressing HEK293T cells. Gray lines represent linear fits on individual cells. Black line is the average fit across all cells (n=9). (**E**) Representative lifetime images of CAESR in HEK293T cells. Scale bars represent 20 μm.

### Figure 2 Supplements

**Fig. 2, S1.**
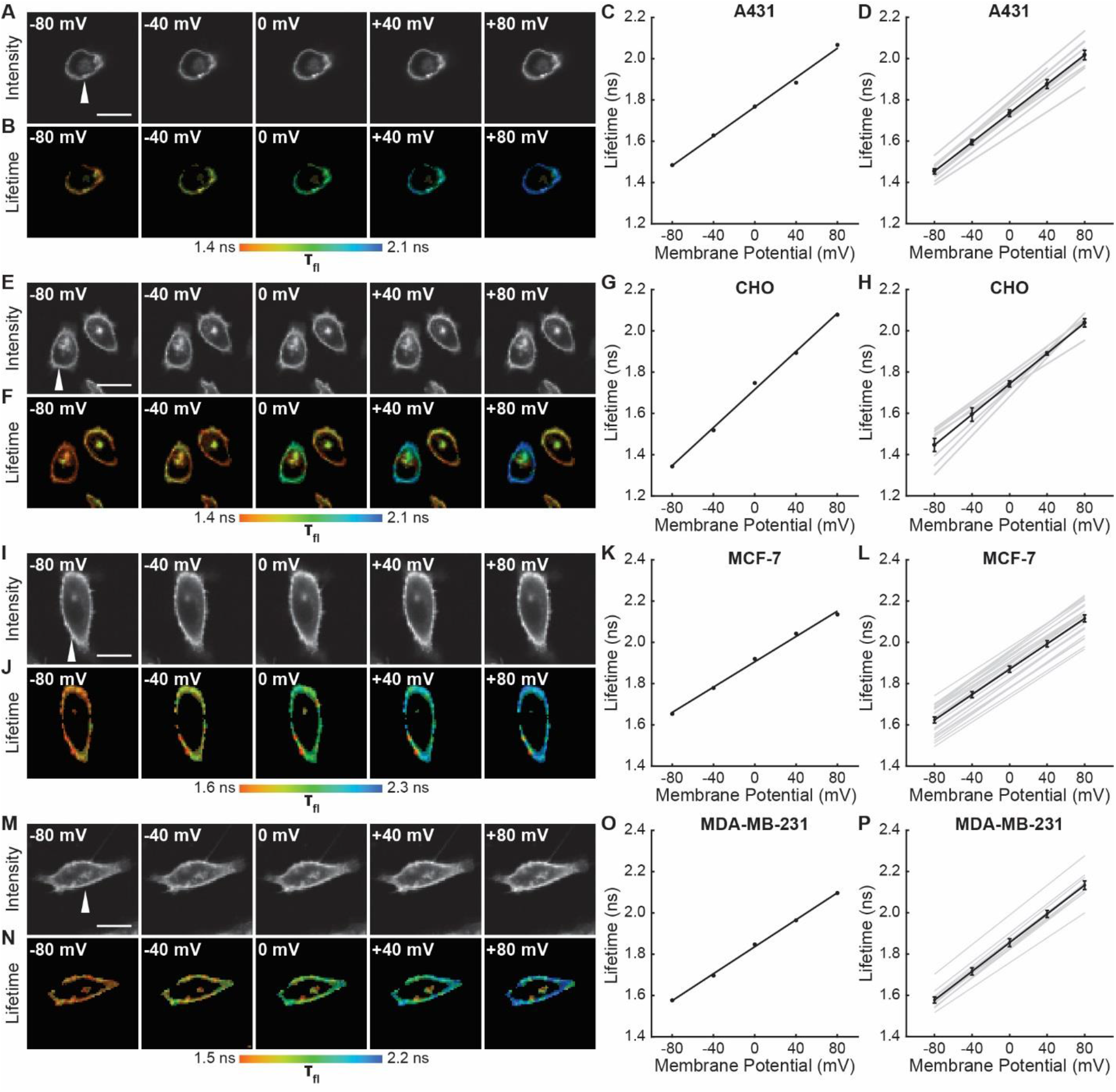
VoltageFluor lifetime reports voltage in various cell types. (**A**) Representative photon count and (**B**) lifetime images of a VF2.1.Cl in A431 cells with V_mem_ held at the indicated value with voltage-clamp electrophysiology. A431 cells were not serum starved for these experiments. (**C**) Quantification of the images in (B), with the line of best fit for this single trial. (**D**) Lines of best fit for the lifetime-V_mem_ relationships of 12 A431 cells (gray lines). Average lifetime at each potential is shown as mean ± SEM, with the average line of best fit in black. (**E**)-(**H**) Lifetime-V_mem_ standard curve determination in CHO cells (n=8). (**I**)-(**L**) Lifetime-V_mem_ standard curve determination in MCF-7 cells (n=24). (**M**)-(**P**) Lifetime-V_mem_ standard curve determination in MDA-MB-231 cells (n=11). VF2.1.Cl concentration was 100 nM in all cases. White arrows indicates the voltage-clamped cell. Scale bars are 20 μm.

**Fig. 2, S2.**
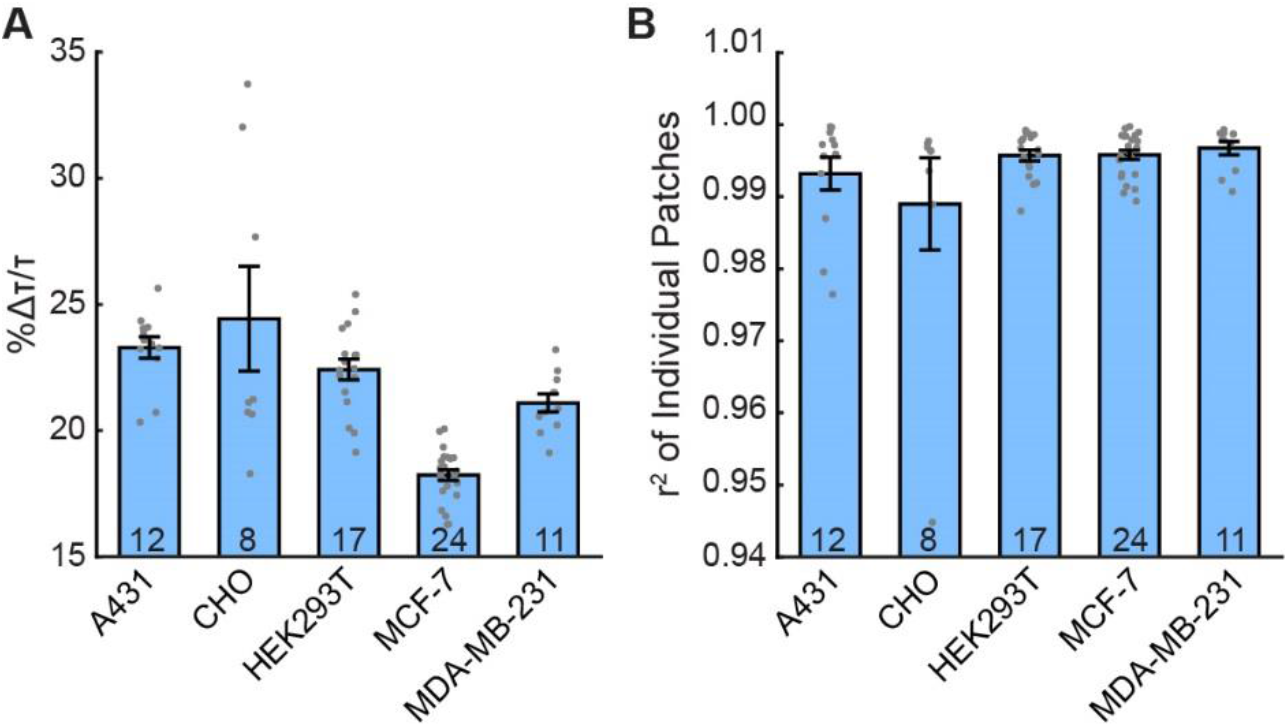
Additional parameters of lifetime-voltage standard curves. (**A**) Percent change in VF2.1.Cl lifetime per 100 mV change in voltage, relative to the lifetime at −60 mV. (**B**) Correlation coefficients (r^2^) for all of the lines of best fit of VF2.1.Cl lifetime versus membrane potential. Values shown are mean ± S.E.M., with gray dots indicating values from individual patches.

**Fig. 2, S3.**
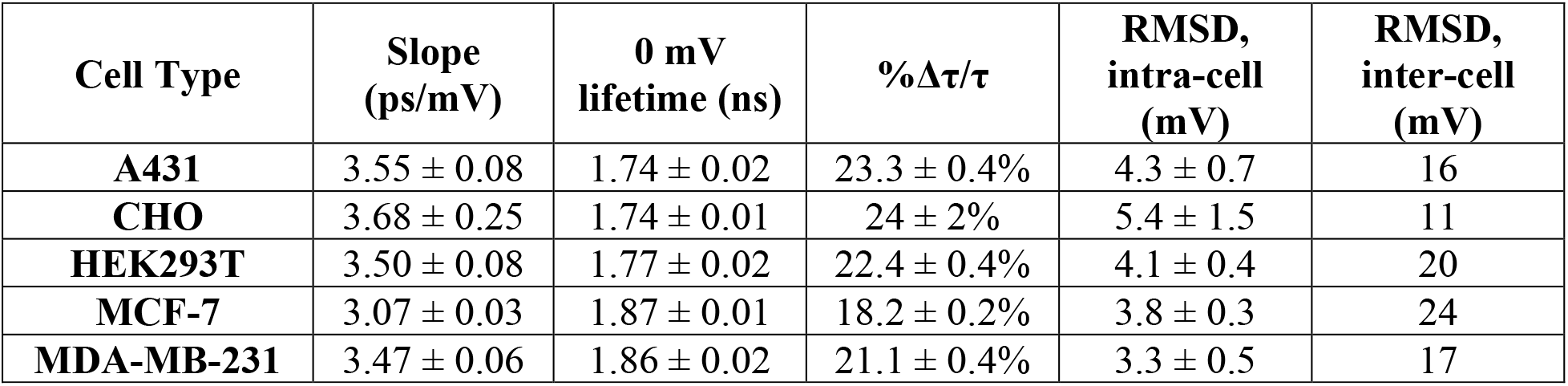
Lifetime-V_mem_ standard curves for VF2.1.Cl lifetime in various cell lines. Whole-cell voltage-clamp electrophysiology was used to determine the relationship between VF2.1.Cl lifetime and membrane potential in five different cell lines. Parameters of this linear model are listed above. The %Δτ/τ is the percent change in the lifetime observed for a voltage step from −60 mV to +40 mV. The intra-cell RMSD represents the accuracy for quantifying voltage changes in a particular cell (see **Methods**). The inter-cell RMSD represents the expected variability in singletrial absolute V_mem_ determinations. Sample sizes: A431 12, CHO 8, HEK293T 17, MCF-7 24, MDA-MB-231 11. All values are tabulated as mean ± SEM.

**Fig. 2, S4.**
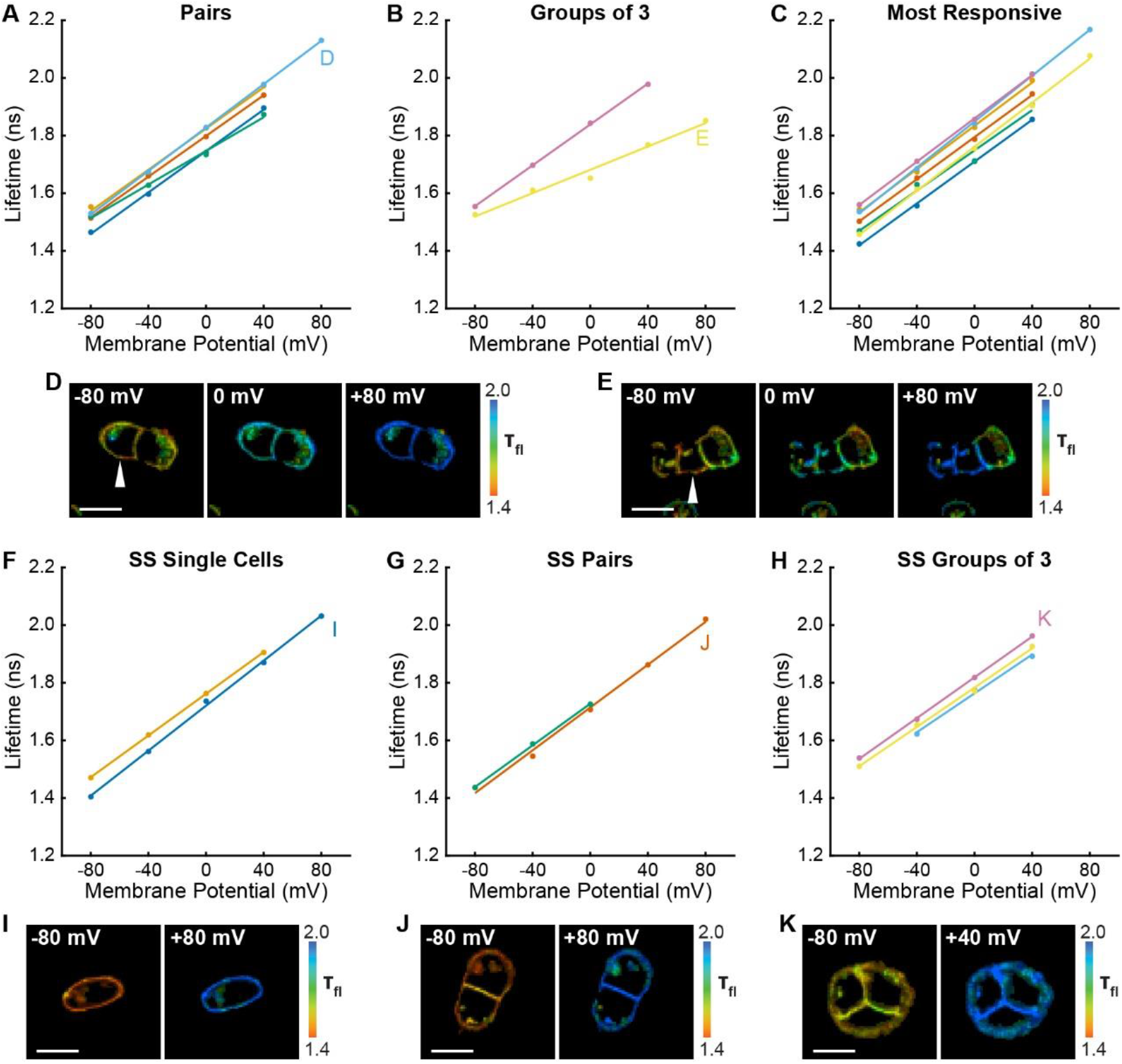
Relationship between lifetime and membrane potential extends to groups of cells and across culture conditions. Electrophysiological calibration of lifetime was performed on small groups of A431 cells and on serum starved (SS) A431 cells to verify that the V_mem_-lifetime standard curves for a given cell line are generalizable across many cellular growth conditions. For all graphs, each line represents a group of cells. Letters on the graphs indicate the subfigure where images from that recording are shown. (**A**) Lifetime-voltage relationships in cell pairs, in which only one cell was directly controlled with voltage-clamp electrophysiology. (**B**) Lifetime-voltage relationships in groups of three cells, in which only one cell was directly controlled with voltage-clamp electrophysiology. (**C**) Lifetime for the most responsive cell from pairs and groups of three in (A) and (B). Line color codes are maintained from (A) and (B). (**D, E**) Representative lifetime images from (A) and (B) respectively. White arrow indicates cell directly controlled with electrophysiology. (**F**) Lifetime-voltage relationship in SS single cells, (**G**) pairs, and (**H**) groups of three cells. (**I**)-(**K**) Representative images from (F)-(H). Scale bars are 20 μm.

**Fig. 2, S5.**
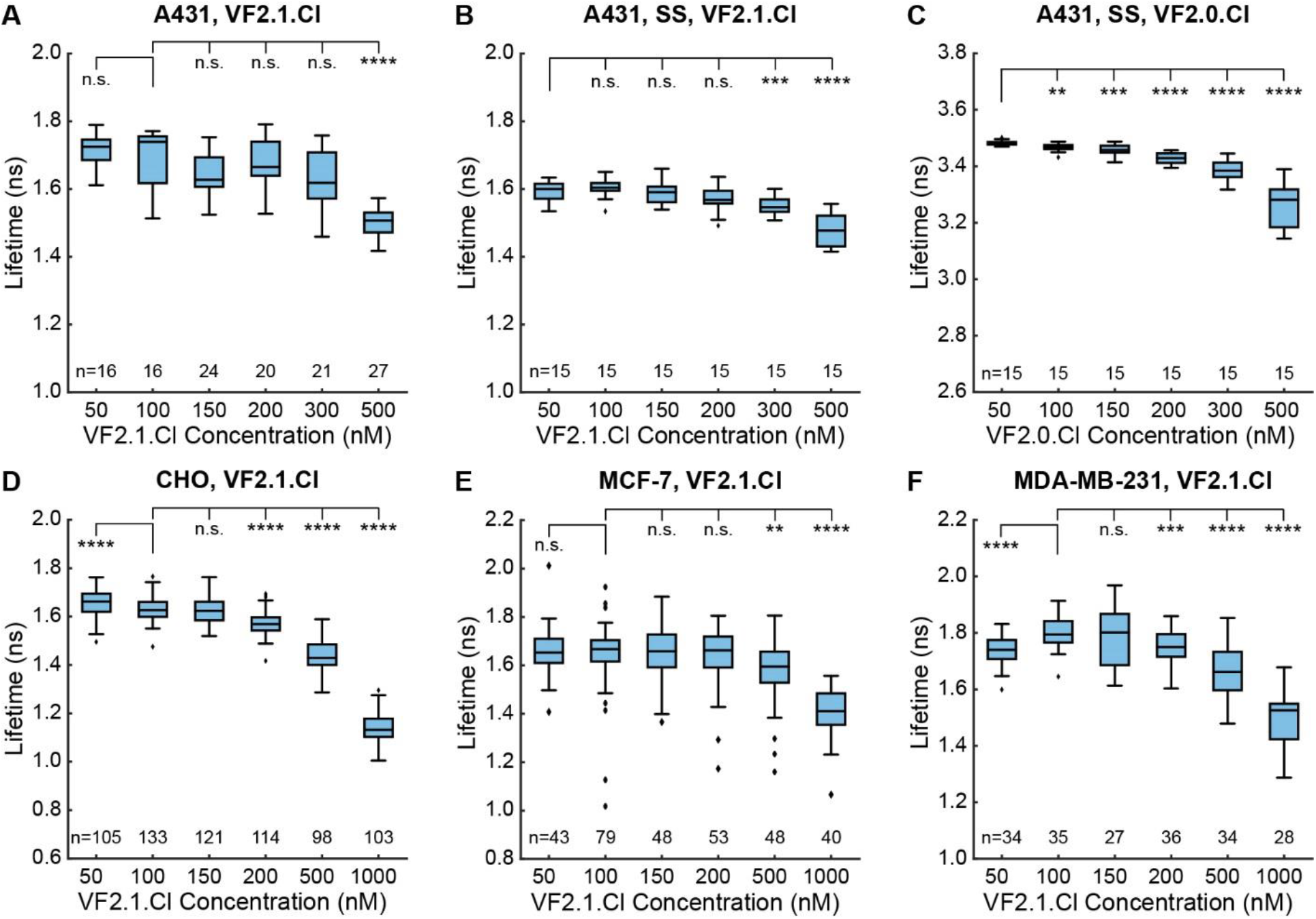
Concentration dependence of VoltageFluor lifetime in four cell lines. A431 cells were analyzed with VF2.1.Cl both in (**A**) full serum and (**B**) serum-starved conditions. (**C**) VF2.0.Cl in serum-starved A431 cells. (**D**) VF2.1.Cl in CHO cells. (**E**) VF2.1.Cl in MCF-7 cells. (**F**) VF2.1.Cl in MDA-MB-231 cells. All VF2.1.Cl data were fit with a biexponential model, and all VF2.0.Cl data were fit with a monoexponential model. Box plots represent the interquartile range, with whiskers and outliers determined with the Tukey method. Sample sizes indicate number of cell groups. Data were acquired over 2 to 4 different days from a total of 3 or 4 coverslips at each concentration. Asterisks indicate significant differences between the indicated concentration and the VF concentration selected for additional experiments (n.s. p > 0.05, * p < 0.05, ** p < 0.01, *** p < 0.001, **** p < 0.0001, two-sided, unpaired, unequal variances t-test).

### Figure 3 Supplements

**Fig. 3, S1.**
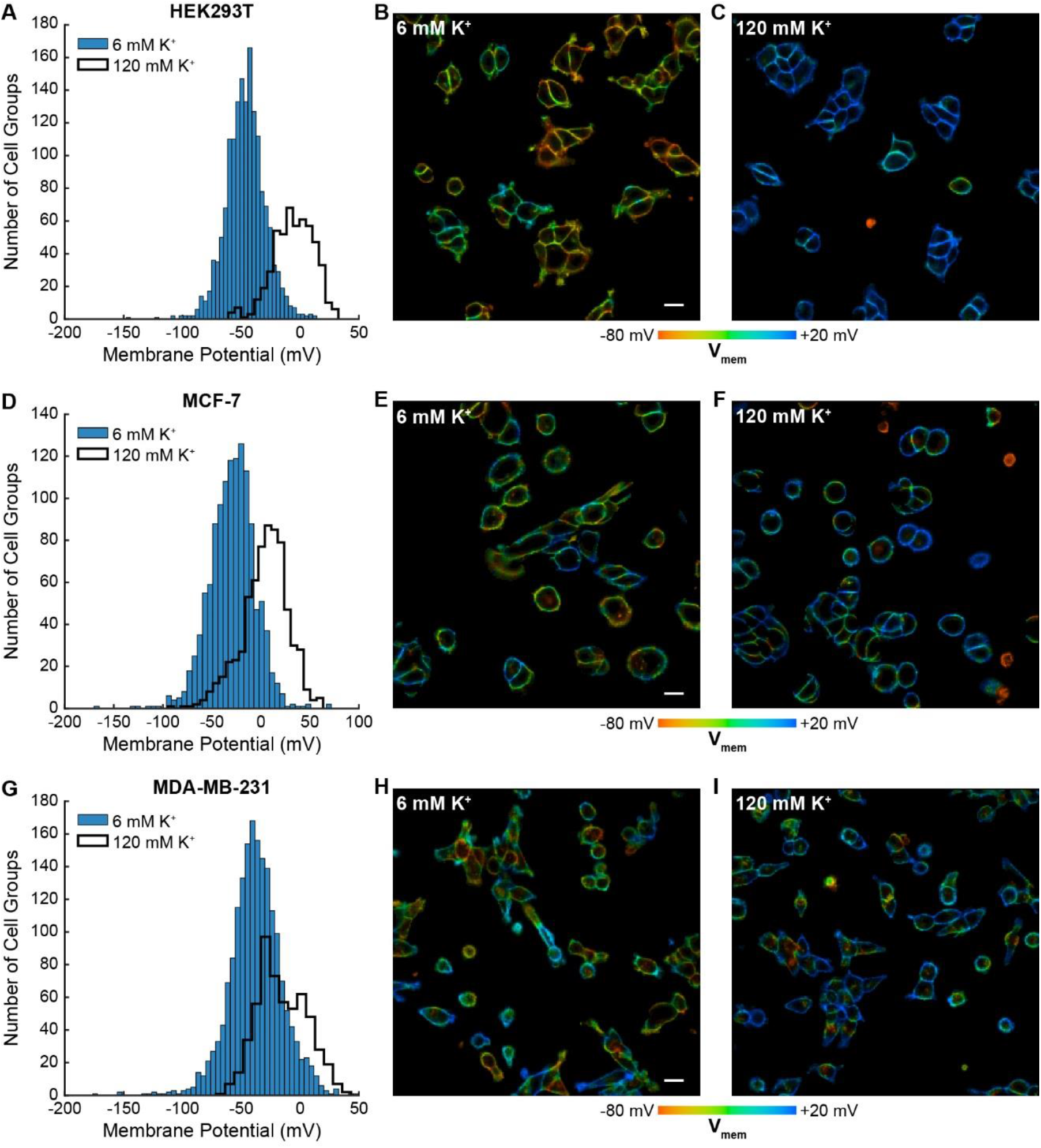
Optically recorded V_mem_ distributions in HEK293T, MCF-7 and MDA-MB-231 cells. Fluorescence lifetime images of cells incubated with 100 nM VF2.1.Cl were used to determine V_mem_ from previously performed electrophysiological calibration (**Fig. 2**). (**A**) Histograms of V_mem_ values recorded in HEK293T cells incubated with 6 mM extracellular K^+^ (commercial HBSS, n=1613) or 120 mM K^+^ (high K^+^ HBSS, n=520). (**B**) Representative lifetime image of HEK293T cells with 6 mM extracellular K^+^. (**C**) Representative lifetime image of HEK293T cells in 120 mM extracellular K^+^. (**D**) Histograms of V_mem_ values observed in MCF-7 cells under normal (n=1259) and high K^+^ (n=681) conditions. Representative lifetime images of MCF-7 cells in (**E**) 6 mM and (**F**) 120 mM extracellular K^+^. (**G**) Histograms of V_mem_ values observed in MDA-MB-231 cells under normal (n=1840) and high K^+^ (n=558) conditions. Representative lifetime images of MDA-MB-231 cells in (**H**) 6 mM and (**I**) 120 mM extracellular K^+^. Histogram bin sizes were determined by the Freedman-Diaconis rule. Intensities in the lifetime-intensity overlay images are not scaled to each other. Scale bars, 20 μm.

**Fig. 3, S2.**
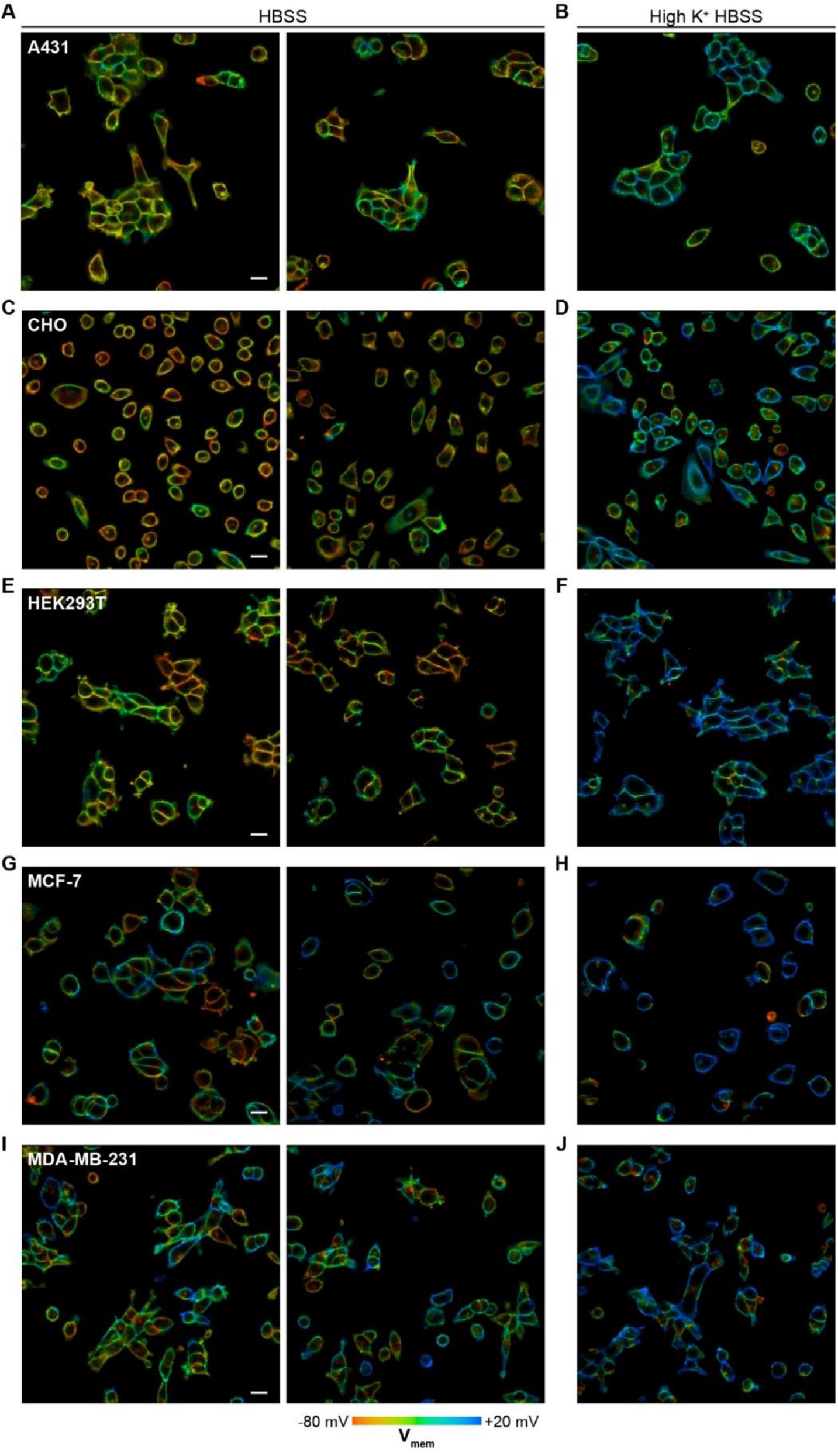
Representative images of cultured cell resting membrane potential. Representative VF-FLIM images of cells in standard imaging buffer (HBSS, 6 mM extracellular K_+_) and high K_+_ imaging buffer (high K_+_ HBSS, 120 mM extracellular K_+_). Membrane potential was calculated per cell group; analyses of pixel by pixel differences in lifetime fall beyond the resolution limit of the VF-FLIM calibrations in this work. Images depict A431 cells in (**A**) HBSS and (**B**) high K^+^ HBSS; CHO cells in (**C**) HBSS and (**D**) high K^+^ HBSS; HEK293T cells in (**E**) HBSS and (**F**) high K^+^ HBSS; MCF-7 cells in (**G**) HBSS and (**H**) high K^+^ HBSS, and MDA-MB-231 cells in (**I**) HBSS and (**J**) high K^+^ HBSS.

**Fig. 3, S3.**
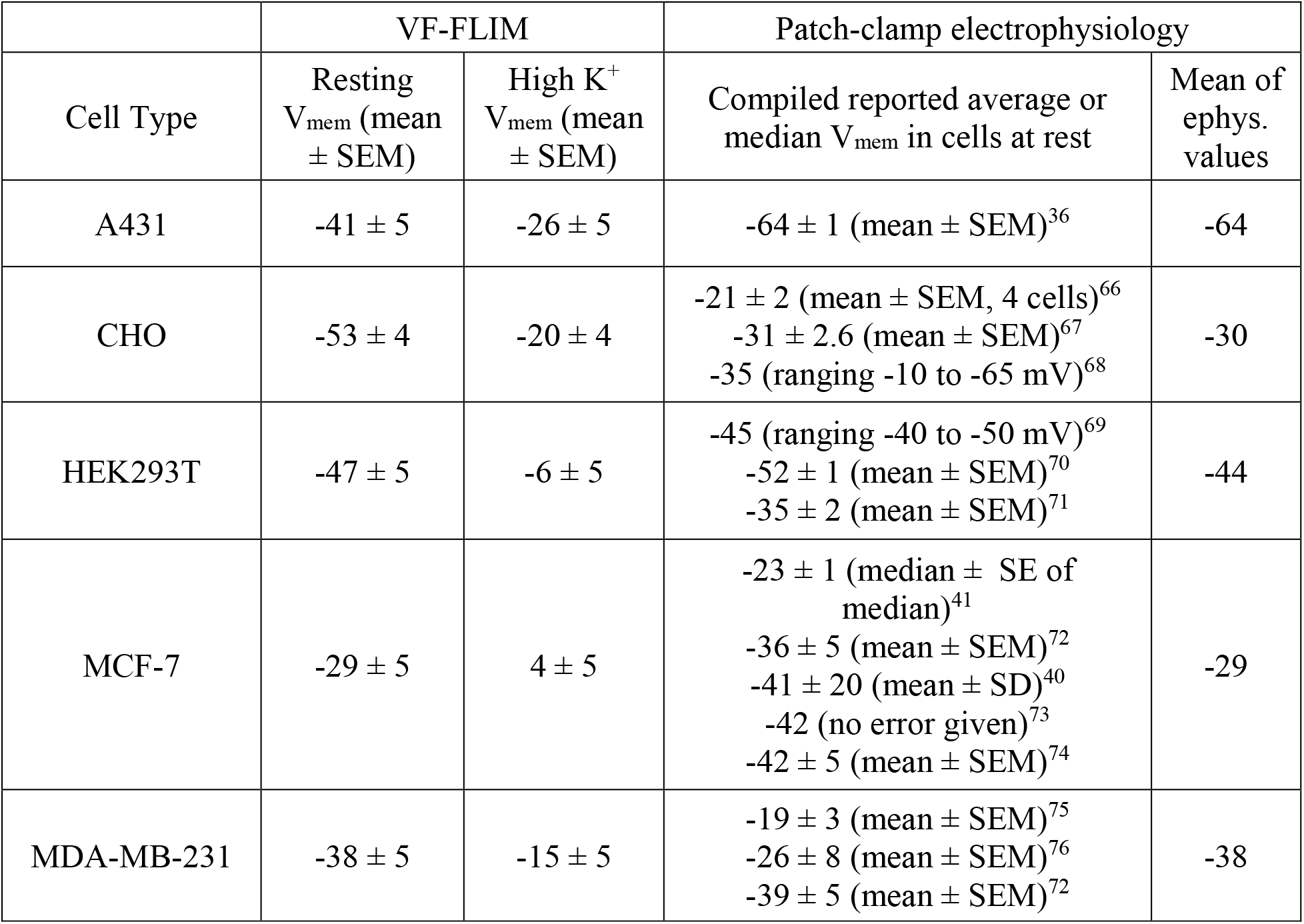
V_mem_ measurements made with VF-FLIM agree with previously reported values. Comparison of optically-determined resting membrane potential values (in millivolts) and previously reported values. This table summarizes data presented in **Fig. 3** and **Fig. 3 – supplement 1**. Optically determined membrane potentials were calculated from lifetime-V_mem_ standard curves (**Fig. 2 – supplement 3**). For tabulated literature values, measures of error and central tendency were used from the original publication. In some cases, none were given or only ranges were discussed. The mean of the reported ephys values is the mean of the values listed here. Sample sizes for resting and elevated K^+^, respectively: A431 1056, 368; CHO 2410, 1310; HEK293T 1613, 520; MCF-7 1259, 681; MDA-MB-231 1840, 558.

### Figure 4 Supplements

**Fig. 4, S1.**
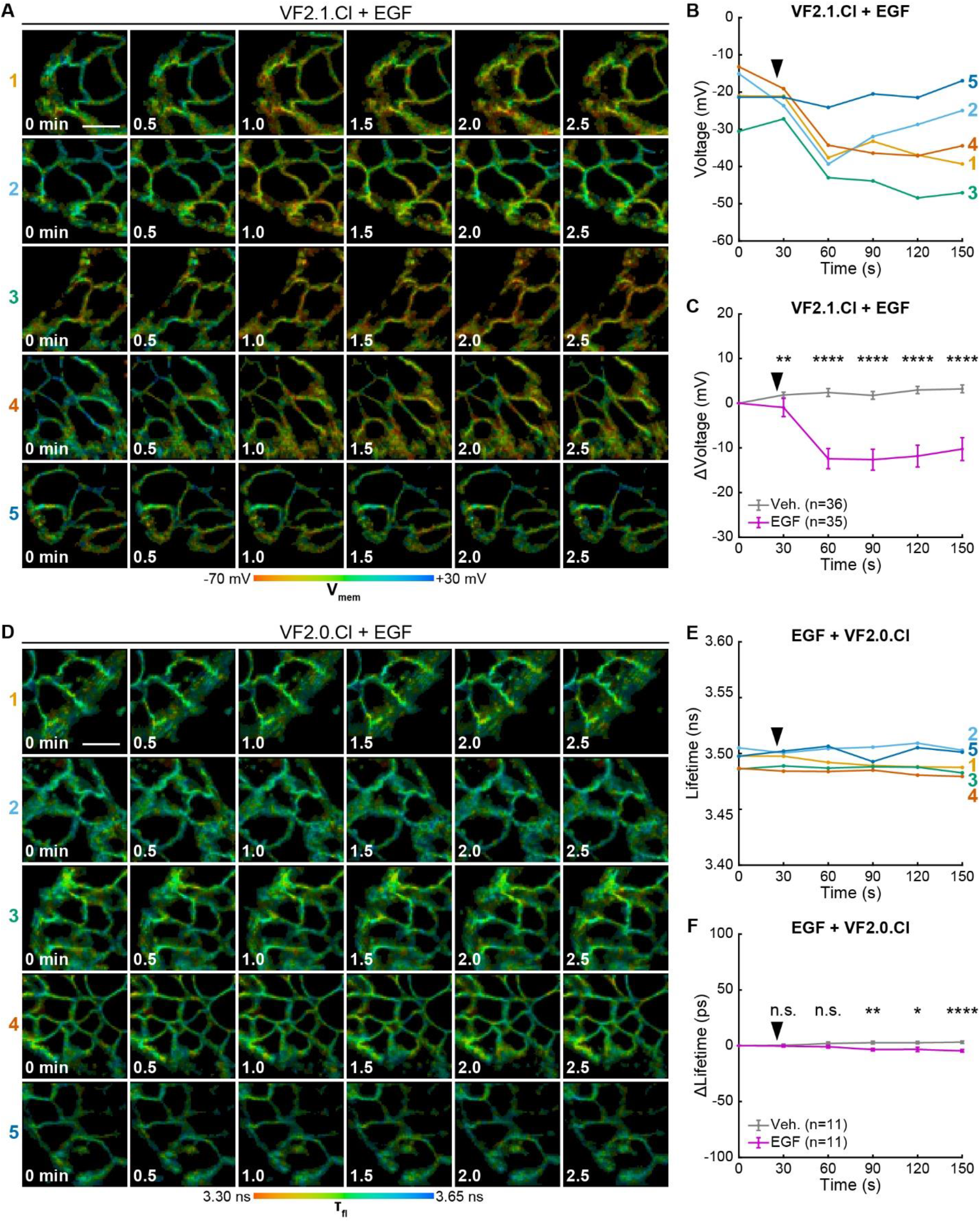
Individual VF-FLIM recordings of A431 EGF response. (**A**) Representative 3 minute VF-FLIM recordings of A431 cells loaded with 50 nM VF2.1.Cl. 500 ng/mL EGF was added 30 seconds into the time series (black arrow). (**B**) Quantification of the images in (A), with a single trace per image series shown. (**C**) Average voltage change in A431 cells following the addition of imaging buffer vehicle (gray) or EGF (purple). (**D**) Control VF2.0.Cl (not voltage sensitive, 50 nM) images of A431 cells treated as in (A). Images are scaled across the same amount of lifetime space (350 ps) as the VF2.1.Cl images. (**E**) Quantification of the images in (D). (**F**) Average VF2.0.Cl lifetime change seen in A431 cells following the addition of imaging buffer vehicle (gray) or EGF (purple) in A431 cells. Graph is scaled across the same amount of lifetime space as the VF2.1.Cl data in (C). Asterisks indicate significant differences between vehicle and EGF treated cells at a given time point (n.s. p > 0.05, * p < 0.05, ** p < 0.01, *** p < 0.001, **** p < 0.0001, two-sided, unpaired, unequal variances t-test). Scale bars are 20 μm.

**Fig. 4, S2.**
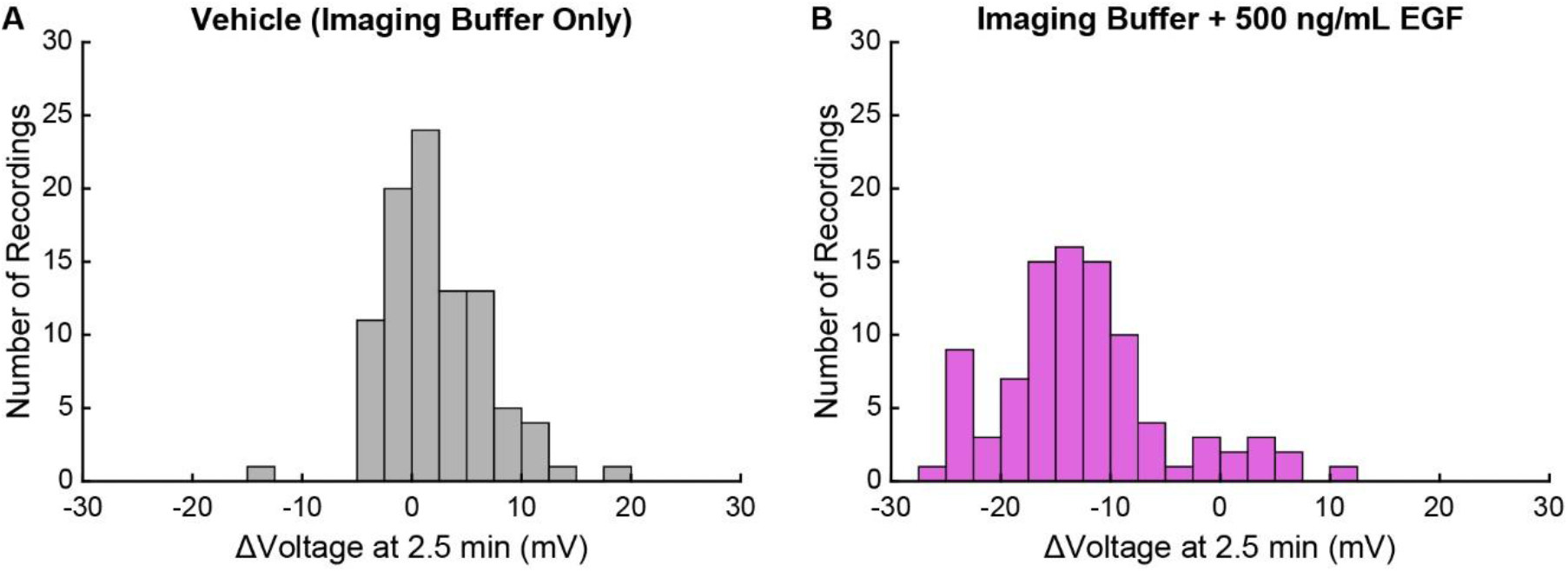
Membrane potential changes in A431 cells 2.5 minutes after EGF treatment. Comparison of V_mem_ changes observed in A431 cells 2.5 minutes after treatment with (**A**) imaging buffer vehicle or (**B**) 500 ng/mL EGF. Data shown here are compiled from **Fig. 4C** and **Fig. 5A** to provide a sense of overall distribution of the responses. Each recording contained a single group of approximately 5 to 10 cells. Sample sizes (number of recordings): Vehicle 93, EGF 92.

**Fig. 4, S3.**
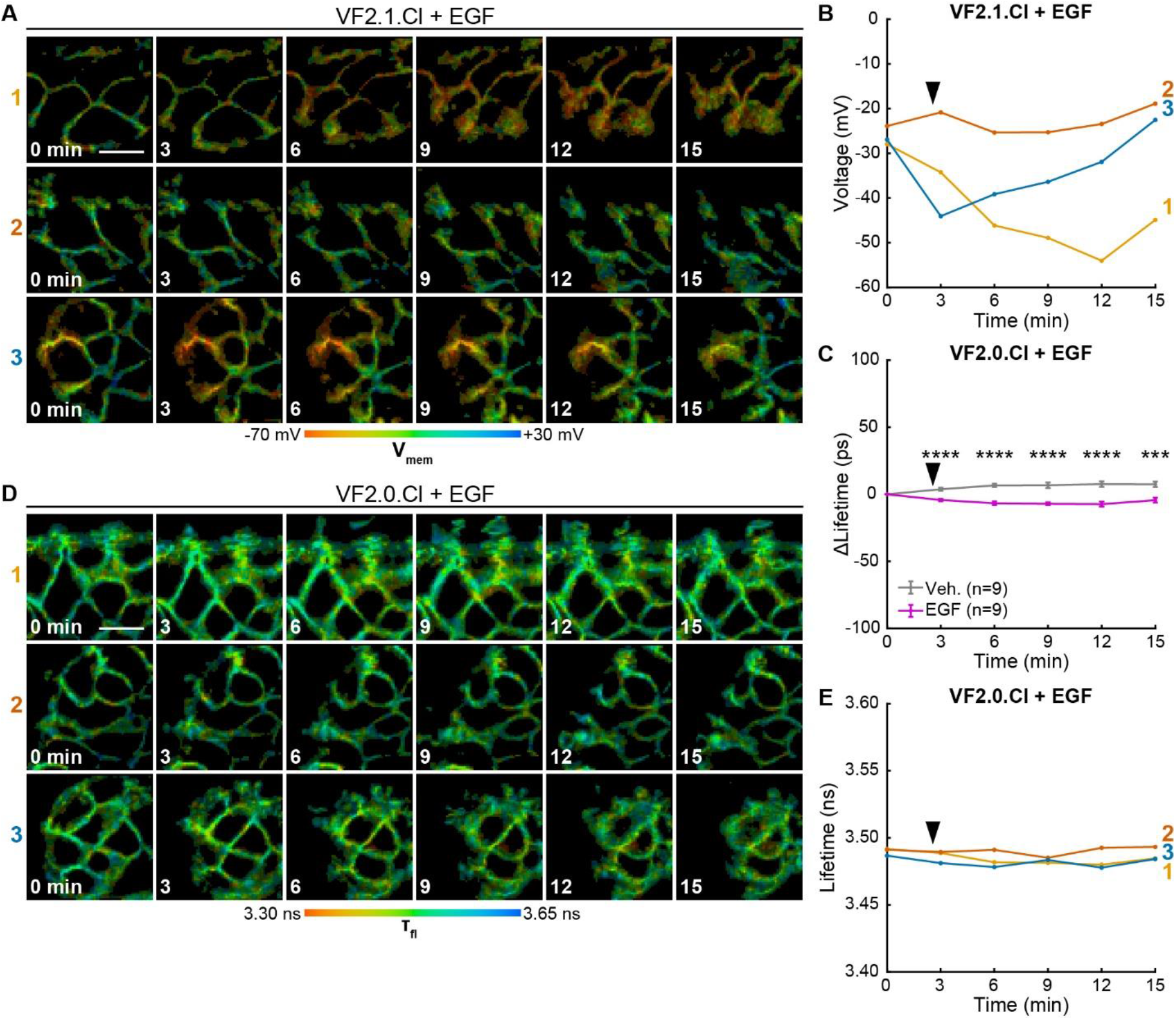
VF-FLIM reports A431 V_mem_ changes over 15 minutes. (**A**) Representative longer term VF-FLIM recordings of A431 cells loaded with 50 nM VF2.1.Cl. 500 ng/mL EGF was added 30 seconds into the time series. (**B**) Quantification of the images in (A), with a single trace per image series shown. (**C**) Control VF2.0.Cl (not voltage sensitive, 50 nM) images of A431 cells treated as in (A). Images are scaled across the same total lifetime range (350 ps) as the VF2.1.Cl images. (**D**) Quantification of the recordings in (C). (E) Average VF2.0.Cl lifetime change seen in A431 cells following the addition of imaging buffer vehicle (gray) or EGF (purple). Asterisks indicate significant differences between vehicle and EGF treated cells at a given time point (*** p < 0.001, **** p < 0.0001, two-sided, unpaired, unequal variances t-test). Scale bars are 20 μm.

**Fig. 4, S4.**
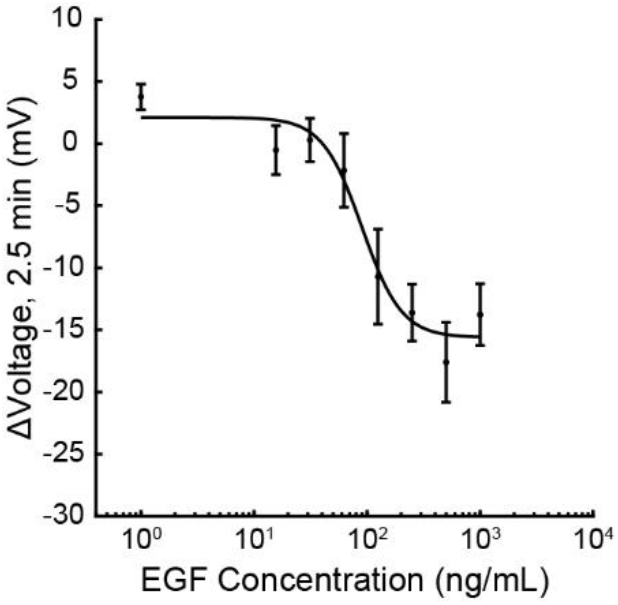
Dose-response relationship of A431 voltage response to EGF. Data were fit to a four-parameter logistic function to obtain an EC50 of 90 ng/mL (95% CI: 47-130 ng/mL). Response to each EGF concentration is shown as mean ± SEM of 6 or 7 recordings (one group of 5-10 cells per recording).

**Fig. 4, S5.**
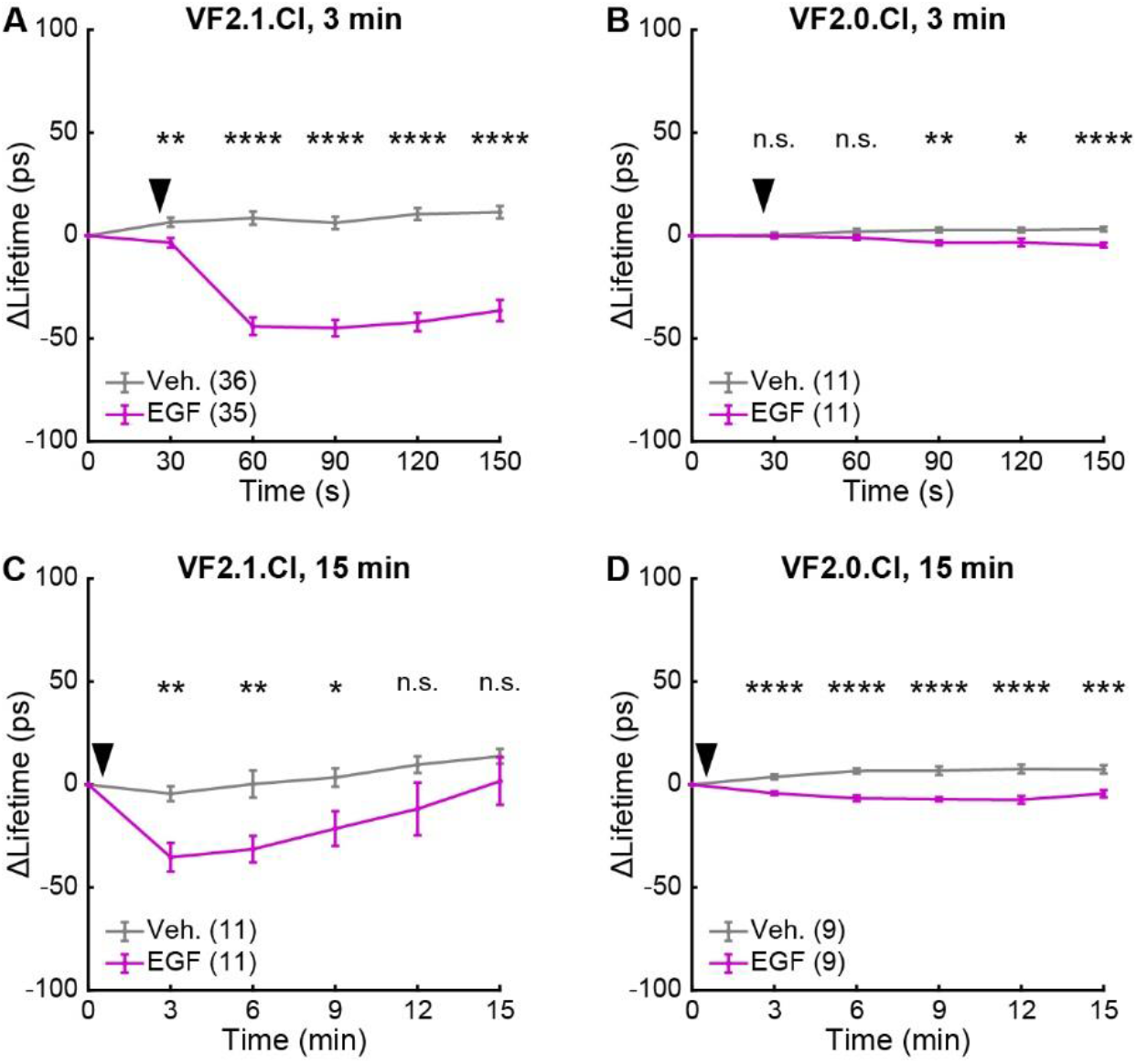
Effect sizes of VF2.1.Cl and VF2.0.Cl response to EGF treatment. Average lifetime changes observed in A431 cells following the addition (black arrow) of imaging buffer vehicle (gray) or 500 ng/mL EGF (purple). (**A**) Cells incubated with 50 nM VF2.1.Cl and imaged for 3 minutes. (**B**) Cells incubated with 50 nM VF2.0.Cl (not voltage sensitive) and imaged for 3 minutes. (**C**) Cells incubated with 50 nM VF2.1.Cl and imaged intermittently for 15 minutes. (**D**) Cells incubated with 50 nM VF2.0.Cl (not voltage sensitive) and imaged intermittently for 15 minutes. Data are reproduced from **Fig. 4, Fig. 4-supplement 1**, and **Fig. 4 - supplement 3**, but here data are scaled in units of lifetime rather than voltage for facile comparison. Data are shown as mean ± SEM for the indicated number of recordings (one group of 5-10 cells per recording). Asterisks indicate significant differences between vehicle and EGF treated cells at a given time point (n.s. p > 0.05, * p < 0.05, ** p < 0.01, *** p < 0.001, **** p < 0.0001, two-sided, unpaired, unequal variances t-test).

**Fig. 4, S6.**
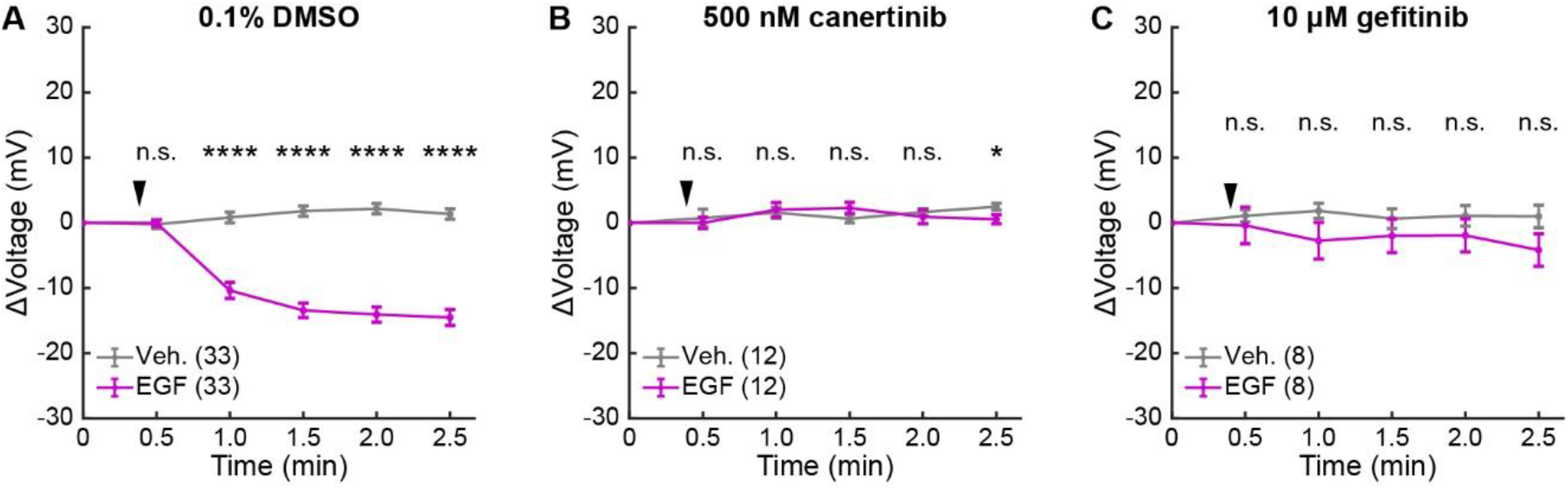
EGFR inhibitors abolish voltage response to EGF in A431 cells. Average V_mem_ changes following the addition (black arrow) of imaging buffer vehicle (gray) or 500 ng/mL EGF (purple) to A431 cells pre-treated with the indicated drug or DMSO vehicle. 2.5 minute time points from this data are shown elsewhere (**Fig. 4J**); entire time series are shown here. Data are presented as mean ± SEM for the indicated number of recordings (one group of 5-10 cells per recording). Asterisks indicate significant differences between vehicle and EGF treated cells at a given time point (n.s. p > 0.05, * p < 0.05, ** p < 0.01, *** p < 0.001, **** p < 0.0001, two-sided, unpaired, unequal variances t-test).

### Figure 5 Supplements

**Fig. 5, S1.**
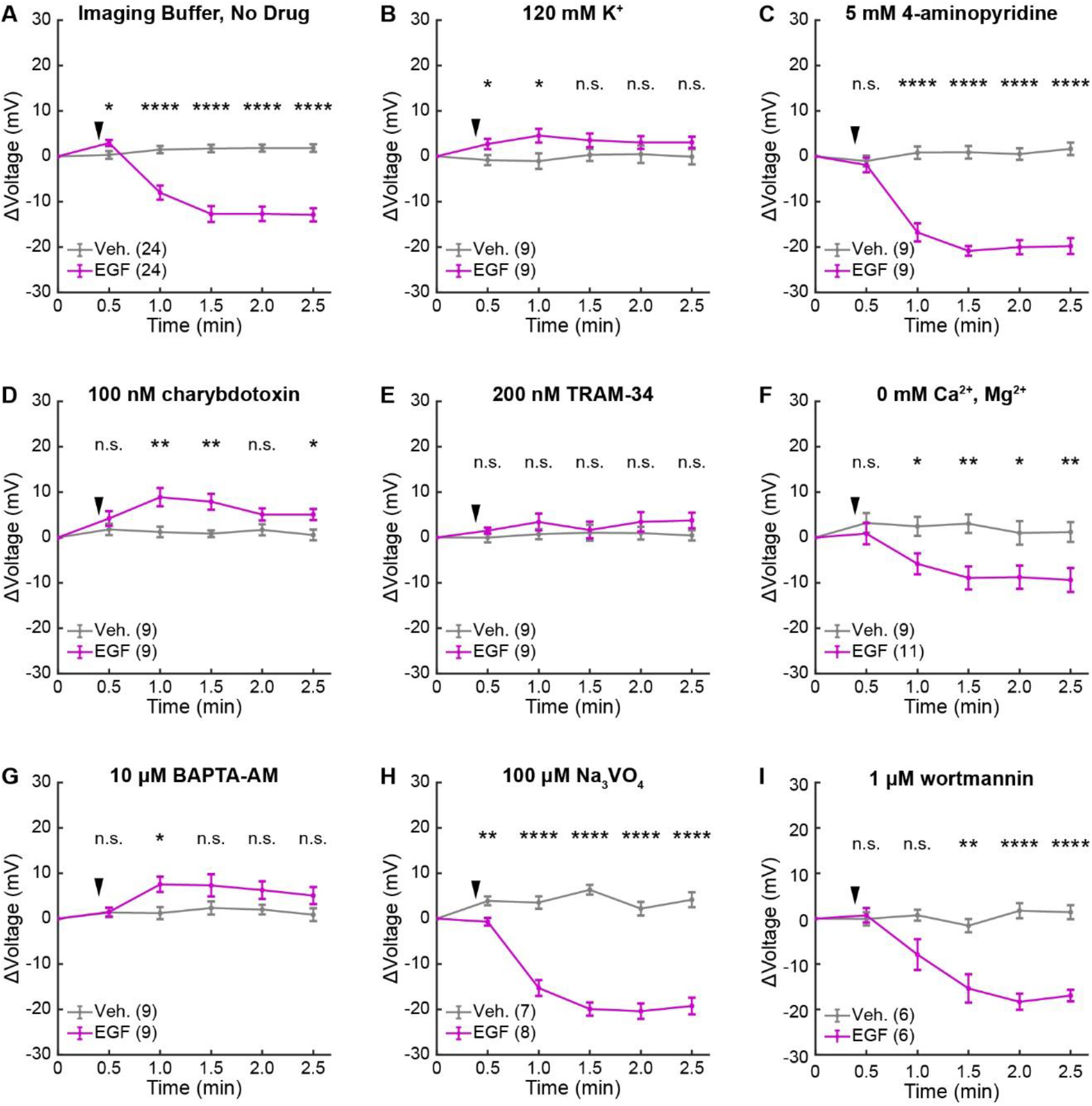
A431 voltage response to EGF with pharmacological intervention. Average V_mem_ changes following the addition (black arrow) of imaging buffer vehicle (gray) or 500 ng/mL EGF (purple) to A431 cells pre-treated with the indicated drug or ionic composition change. 2.5 minute time points from this data are shown elsewhere (**Fig. 5**); entire time series are shown here to illustrate the time courses of the large hyperpolarizing current and small depolarizing current. Data are shown as mean ± SEM for the indicated number of recordings (one group of 5-10 cells per recording). Asterisks indicate significant differences between vehicle and EGF treated cells at a given time point (n.s. p > 0.05, * p < 0.05, ** p < 0.01, *** p < 0.001, **** p < 0.0001, two-sided, unpaired, unequal variances t-test).

**Fig. 5, S2.**
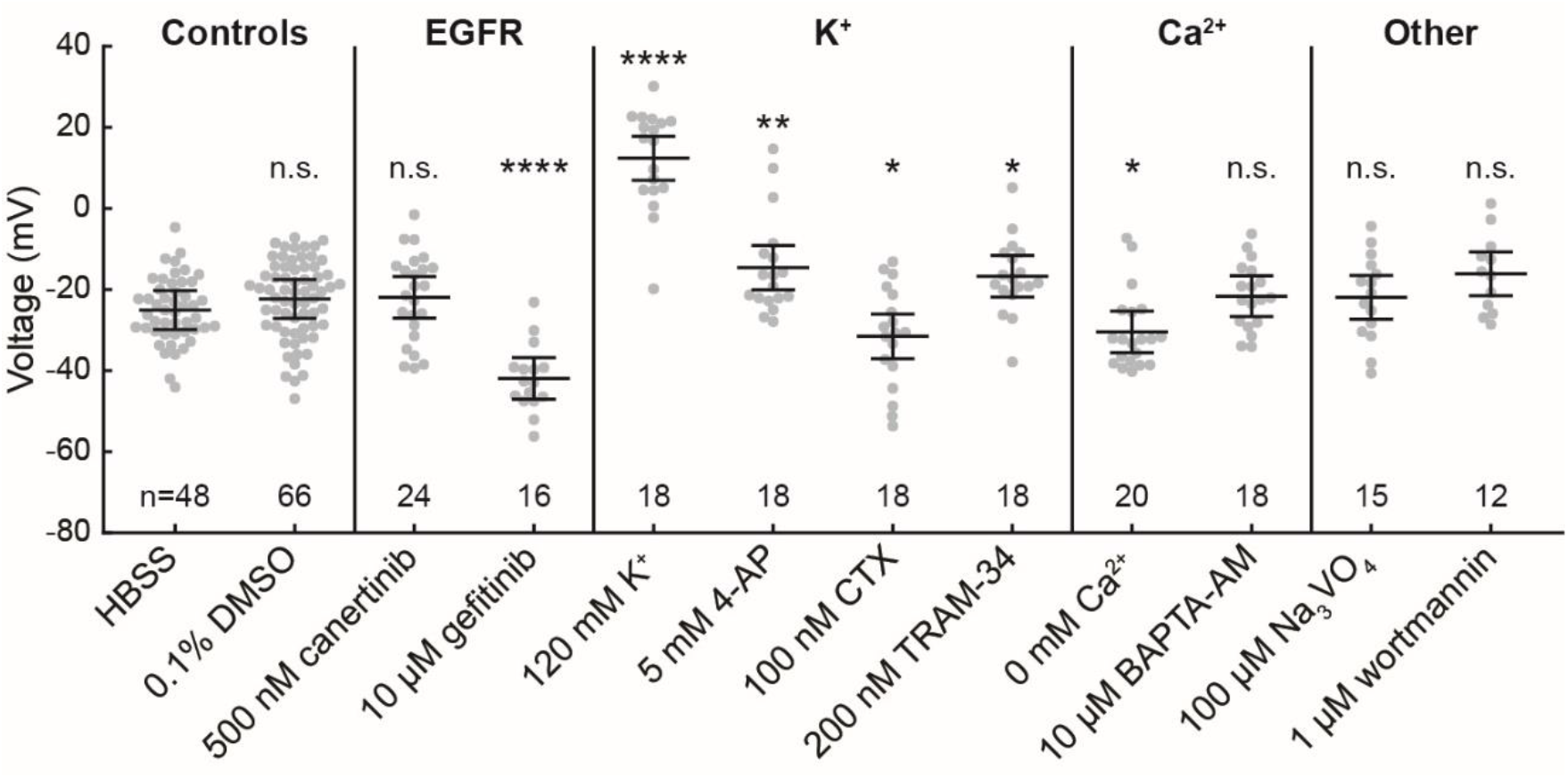
Effects of pharmacological and ionic perturbations on A431 resting membrane potential. Data are the initial V_mem_ reference images for recordings used in EGF addition time series. Data are shown as mean ± SEM for the indicated number of images (one group of 5-10 cells per image), and gray dots represent individual images. Asterisks indicate significant differences between the appropriate vehicle (HBSS or 0.1% DMSO) and pharmacology treated cells (n.s. p > 0.05, * p < 0.05, ** p < 0.01, *** p < 0.001, **** p < 0.0001, two-sided, unpaired, unequal variances t-test). CTX = charybdotoxin, 4-AP = 4-aminopyridine, BAPTA-AM = bisaminophenoxyethanetetraacetic acid acetoxymethyl ester.

